# Sex-Dependent Effects of Glutamatergic Disruption on Dopaminergic Neuron Subtype Vulnerable in Parkinson’s Disease

**DOI:** 10.64898/2026.05.06.723291

**Authors:** Kathleen F. Carmichael, Victor M. Martinez Smith, Jinhui Ding, Gavin Riccobono, Lisa Chang, Lixin Sun, Lupeng Wang, Huaibin Cai

**Affiliations:** Transgenic Section, Laboratory of Neurogenetics, National Institute on Aging, National Institutes of Health, Bethesda, MD 20892, U.S.A.; The Graduate Partnership Program of NIH and Brown University, National Institutes of Health, Bethesda, MD 20892, U.S.A.; Computational Biology Group, Laboratory of Neurogenetics, National Institute on Aging, National Institutes of Health, Bethesda, MD 20892, U.S.A

## Abstract

Aldehyde dehydrogenase 1A1-positive (ALDH1A1^+^) dopaminergic neurons (DANs) are preferentially vulnerable in Parkinson’s disease (PD), yet how their activity is modulated by presynaptic inputs remains poorly defined. Here we investigated the role of glutamatergic input by conditionally deleting *Grin1*, which encodes a critical NMDA receptor (NMDAR) subunit, in ALDH1A1^+^ DANs. *Grin1* conditional knockout (cKO) mice displayed normal locomotion and motor learning; however, females exhibited enhanced operant reward acquisition and excessive feeding with transient weight gain following food restriction. To determine regional contributions, *Grin1* was selectively knocked down in ALDH1A1^+^ DANs of either the ventral tegmental area (VTA) or *substantia nigra pars compacta* (SNc). VTA-specific knockdown in females was sufficient to reproduce the post-restriction feeding and weight gain phenotype. Bulk whole-brain mRNA sequencing revealed pronounced sex-dependent transcriptional changes, primarily in female *Grin1* cKO mice after food restriction. Many differentially expressed genes were associated with mitochondrial function, energy metabolism, and synaptic signaling. Together, these findings reveal a sex-specific role for NMDAR–mediated glutamatergic input to ALDH1A1^+^ VTA DANs in regulating feeding behavior, providing mechanistic insight into how dysfunction of this vulnerable subpopulation may contribute to PD-associated compulsive eating disorders.

**Key Finding:** Disrupted NMDA receptor–mediated glutamatergic input to ALDH1A1^+^ DANs drives sex-specific feeding abnormalities relevant to PD-associated compulsive eating disorders.

## Introduction

Parkinson’s disease (PD) is a progressive neurodegenerative disorder marked by motor symptoms such as tremor, rigidity, and bradykinesia, as well as non-motor symptoms including compulsive eating and food addiction [1–3]. A recent case-control study further reported that female PD patients are more likely to develop food addiction [4]. However, the synaptic and cell type–specific mechanisms underlying PD-associated eating disturbances remain poorly understood.

A pathological hallmark of PD is the pronounced degeneration of nigrostriatal dopaminergic neurons (DANs), particularly within the ventral *substantia nigra pars compacta* (SNc) [5, 6]. Among these, a vulnerable subpopulation expresses aldehyde dehydrogenase 1A1 (ALDH1A1) [7], an enzyme that detoxifies reactive dopamine metabolites such as 3,4-dihydroxyphenylacetaldehyde (DOPAL) and contributes to retinoic acid and GABA metabolism [8–10]. ALDH1A1 expression is markedly reduced in postmortem PD brains [7, 11], and ALDH1A1^+^ DANs in the ventral tier of the SNc are preferentially lost in patients [7, 12, 13]. These neurons are conserved across species and implicated in motor control and motor skill learning [14], yet their regulation by afferent inputs in motor and non-motor functions are not fully elucidated.

Glutamatergic input provided the primary excitatory drive onto midbrain DANs and is critical for synaptic plasticity underlying learning and motivated behavior [15, 16]. N-methyl-D-aspartate receptors (NMDARs), a subtype of ionotropic glutamate receptor, regulate burst firing, a defining electrophysiological feature of ALDH1A1^+^ DANs [17, 18]. Previous studies eliminating NMDARs in all DANs from embryonic stages found that basic dopamine-dependent behaviors such as locomotion and operant learning were largely preserved, whereas deficits emerged in habit learning, cue-dependent conditioning, and effort-related decision-making [19–21]. However, these manipulations did not specifically examine ALDH1A1^+^ subpopulations in adult animals or distinguish between the SNc and ventral tegmental area (VTA), which are differentially involved in motor and reward processing.

Here, we selectively disrupted NMDAR-mediated glutamatergic signaling in ALDH1A1^+^ DANs of the SNc and VTA in adult mice to examine their role in locomotion, motor skill learning, motivation, and feeding behavior. Given the preferential vulnerability of ALDH1A1^+^ DANs in PD-related neurodegeneration [22, 23], this approach enables direct investigation of how excitatory regulation of this subpopulation contributes to PD-relevant motor and non-motor phenotypes. To our knowledge, this is the first study to define the sex-specific behavioral and transcriptomic consequences of disrupting glutamatergic input to ALDH1A1^+^ DANs.

## Results

### Genetic deletion of *Grin1* in adult ALDH1A1^+^ midbrain DANs

To selectively disrupt NMDA receptor (NMDAR) signaling, we targeted the Glutamate Ionotropic Receptor NMDA Type Subunit 1 (*Grin1*) gene, which encodes the obligatory GluN1 subunit (also referred to as NR1, NMDAR1, or GluRξ1) [24]. Functional NMDARs require two GluN1 subunits in combination with GluN2 and/or GluN3 subunits [25–30]. Thus, loss of *Grin1* is sufficient to abolish NMDAR-mediated signaling.

We restricted this disruption to ALDH1A1^+^ midbrain DANs by crossing *Aldh1a1*^+/P2A-CreERT2^ mice [14] with floxed *NR1* (*Grin1*^fl/fl^) mice [31]. In the resulting approximately 1–2-month-old *Aldh1a1*^+/P2A-CreERT2^::*Grin1*^fl/fl^ conditional knockout (*Grin1* cKO) mice, 4-hydroxytamoxifen (4-OHT) administration induced Cre-mediated recombination selectively in ALDH1A1-expressing cells, excising the floxed region of *Grin1* (**Fig. 1a**). This inducible strategy provided temporal control, enabling deletion in adulthood to avoid developmental compensation or altered circuit maturation [32–34]. Immunohistochemistry confirmed a marked reduction of GluN1 in ALDH1A1^+^ midbrain DANs of *Grin1* cKO mice, while ALDH1A1-negative neurons retained expression (**Fig. 1b, Supplementary Fig. 1**). No reduction was observed in *Grin1*^fl/fl^ control mice (**Fig. 1b, Supplementary Fig. 1**).

**Fig. 1.**
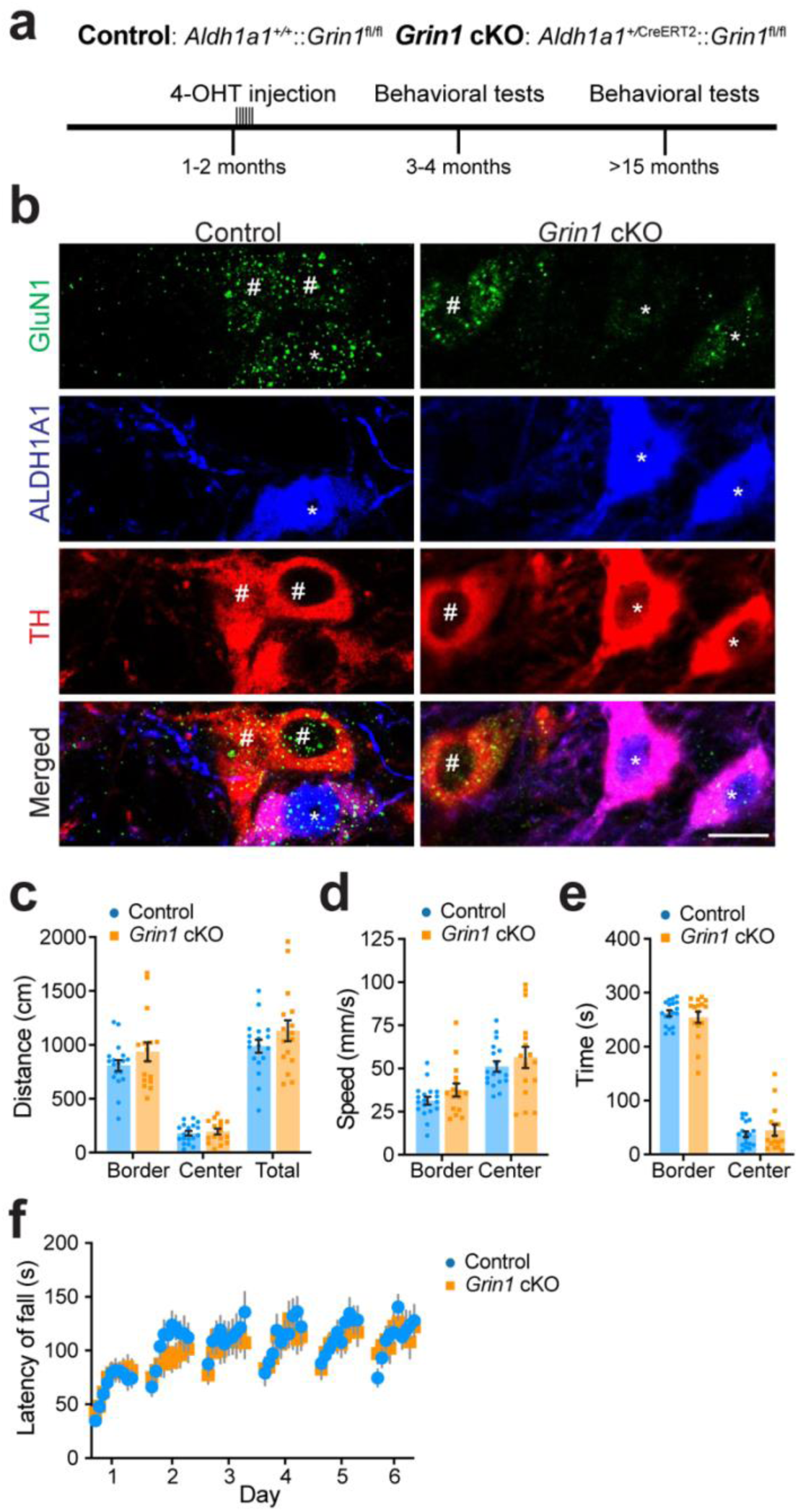
*Grin1* cKO mice exhibit no deficits in spontaneous locomotion or motor skill learning relative to controls. **a,** Experimental timeline showing 4-hydroxytamoxifen (4-OHT) injections and subsequent open field and rotarod testing in two age cohorts following knockout of the *Grin1* in ALDH1A1^+^ midbrain DANs. **b,** Representative confocal images of GluN1 (green), ALDH1A1 (blue), and TH (red) co-staining in midbrain coronal sections from *Aldh1a1*^+/P2A-CreERT2^::*Grin1*^fl/fl^ conditional knockout (*Grin1* cKO) and *Grin1*^fl/fl^ (control) mice (63 ×, single z-plane). TH marks DANs. “*” indicates ALDH1A1^+^ DANs. “#” labels ALDH1A1-negative DANs. Scale bar, 20 μm. **c,** Distance traveled (mm) in the border, center, and total regions of the open field arena by *Grin1* cKO (n = 7 Female [F], 9 Male [M]) and control mice (n = 8F, 10M) during a 5-minute session. Unpaired t-tests with Welch correction: border, p = 0.2160; center, p = 0.6423; total, p = 0.2177. **d,** Average speed (mm/s) in the border and center regions. Unpaired t-tests with Welch correction: border, p = 0.1619; center, p = 0.4448. **e,** Time spent (s) in the border and center regions. Unpaired t-tests with Welch correction: *Grin1* cKO versus control in border, p = 0.5276; *Grin1* cKO versus control in center, p = 0.5315. **f,** Performance on 6-day repeated rotarod tests (10 trials/day) comparing *Grin1* cKO (n = 7F, 10M) and control mice (n = 8F, 12M). Two-way ANOVA, genotype: F(1, 35) = 0.7043, p = 0.7043. All data presented as mean ± SEM.

### Impaired NMDAR-mediated glutamatergic input to ALDH1A1^+^ midbrain DANs does not affect voluntary locomotion or rotarod motor learning

We first assessed whether loss of NMDAR-mediated glutamatergic input to ALDH1A1^+^ midbrain DANs alters spontaneous locomotion using the open field test. In 3–4-month-old *Grin1* cKO and control mice, there were no differences in distance traveled, speed, or time spent in either the border or center of the arena (**Fig. 1c-e**). Sex-stratified analyses confirmed no differences between genotypes (**Supplementary Fig. 2a-f**). Both groups preferred the periphery over the center, consistent with expected anxiety-like behavior [35], indicating that *Grin1* cKO mice did not exhibit altered anxiety. These findings suggest that normal NMDAR-mediated glutamatergic regulation of ALDH1A1^+^ neurons is not required for spontaneous locomotor activity.

Next, we examined motor learning using the accelerating rotarod task (0–40 rpm over 5 min, 10 trials/day for 6 days). Both *Grin1* cKO and control mice improved across trials, but performance did not differ between genotypes (**Fig. 1f**). Sex-specific analyses also revealed no differences (**Supplementary Fig. 2g-h**).

To test whether aging unmasks any potential locomotor or motor learning deficits, we repeated these assays in a separate cohort of ≥15-month-old mice. Aged *Grin1* cKO and control mice showed no differences in open field performance (**Supplementary Fig. 3a-i**) or rotarod learning (**Supplementary Fig. 3j-l**). As observed in young mice, both genotypes improved at the motor learning task over time. Together, these results demonstrate that disrupting NMDAR-mediated glutamatergic input to ALDH1A1^+^ midbrain DANs does not affect spontaneous locomotion or motor learning, and this lack of effect persists with aging.

### Female-specific enhancement in operant learning despite loss of glutamatergic input to ALDH1A1^+^ DANs

To examine effects on non-motor functions, we next tested whether disrupting glutamatergic input to ALDH1A1^+^ midbrain DANs affects operant learning or motivation. Food-restricted mice were trained to lever press for chocolate-flavored pellets (**Fig. 2a**). Each operant chamber contained two levers. Pressing the “active” lever resulted in chocolate pellet reward delivery at the indicated rate reinforcement schedule (except for a 5 second time-out period after pellet delivery), while pressing the “inactive” lever had no consequence. After initial fixed ratio 1 (FR1) training, in which the animal earns one reward for each “active” lever press, mice were challenged with progressively demanding fixed ratio (FR) and progressive ratio (PR) schedules (**Fig. 2a**). Across FR schedules, total pellet acquisition did not differ between genotypes (**Fig. 2b**). When analyzed by sexe, however, female *Grin1* cKO mice consistently earned more pellets than female controls (**Fig. 2c**), whereas no significant differences were observed between male *Grin1* cKO and controls (**Fig. 2d**).

**Fig. 2.**
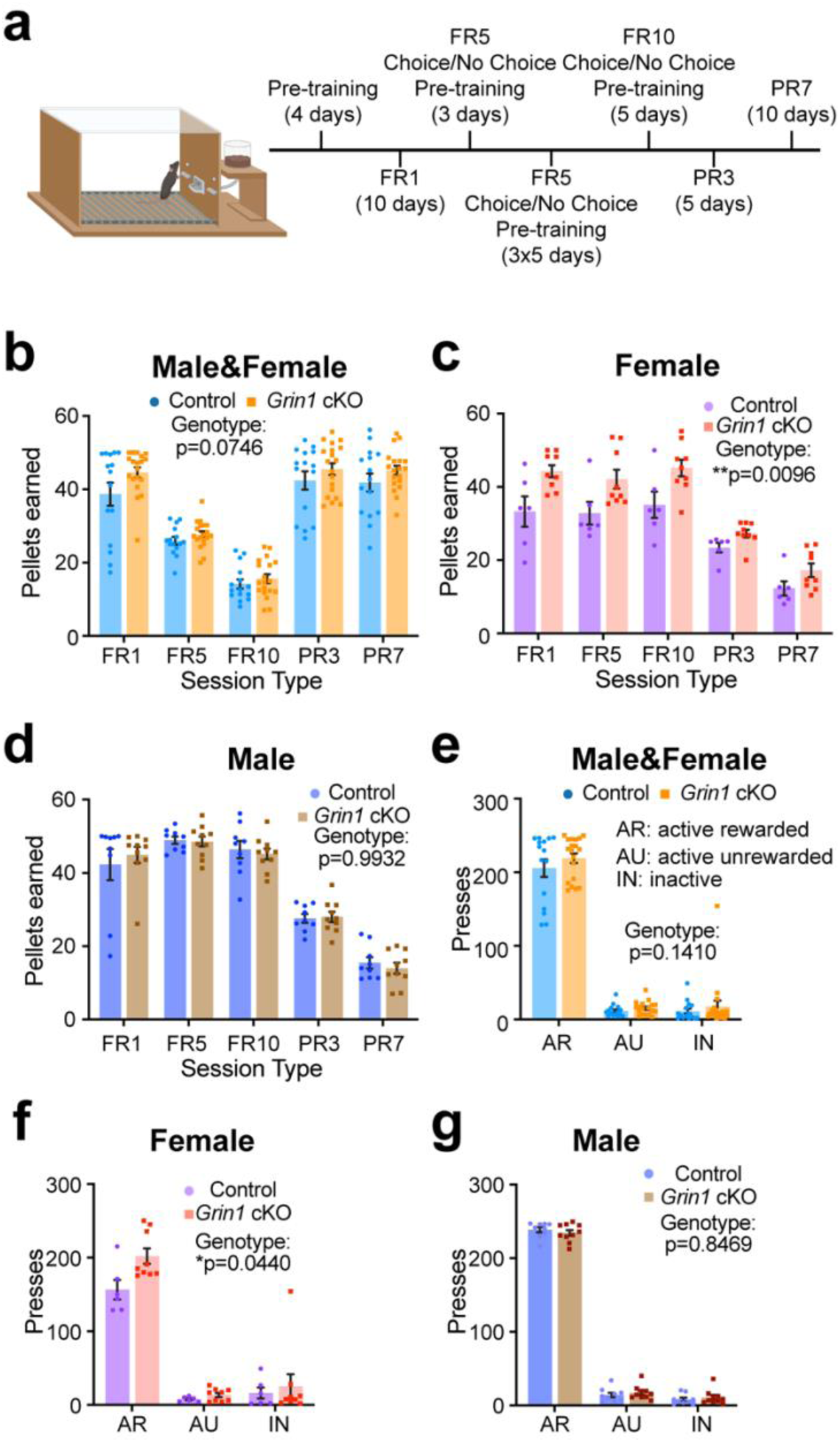
Female *Grin1* cKO mice exhibit sex-specific enhancement in operant learning despite impaired glutamatergic input to ALDH1A1^+^ DANs. a, Schematic of operant chamber setup (created with BioRender) and experimental timeline. **b**, Average number of pellets earned by *Grin1* cKO (n = 9F, 10M) and control mice (n = 6F, 9M) across all reinforcement schedules. Two-way ANOVA, genotype: F(1,32) = 3.396, p = 0.0746. **c**, Average number of pellets earned by female *Grin1* cKO (n = 9F) and control mice (n = 6F). Two-way ANOVA, genotype: F(1,13) = 9.208, **p = 0.0096. Multiple Mann-Whitney tests, Control vs. cKO: FR1, adjusted p = 0.0685; FR5, adjusted *p = 0.0471; FR10, adjusted p = 0.0966; PR3, adjusted *p = 0.0325; PR7, adjusted p = 0.0966. **d**, Average number of pellets earned by male *Grin1* cKO (n = 10M) and control mice (n = 9M). Two-way ANOVA, genotype: F(1,17) = 7.533e-005, p = 0.9932. **e**, Average active and inactive lever press responses by *Grin1* cKO (n = 9F, 10M) and control (n = 6F, 9M) mice during the first ten FR5 sessions. Multiple Mann-Whitney tests, Control vs. cKO: AR, adjusted p = 0.7036; AU, adjusted p = 0.3842; IN, adjusted p = 0.7036. **f**, Average active and inactive lever press responses by female *Grin1* cKO (n = 9F) and control (n = 6F) mice during the first ten FR5 sessions. Multiple Mann-Whitney tests, Control vs. cKO: AR, adjusted *p = 0.0355; AU, adjusted p = 0.5679; IN, adjusted p = 0.8639. **g**, Average active and inactive lever press responses by male *Grin1* cKO (n = 10M) and control (n = 9M) mice during the first ten FR5 sessions. Multiple Mann-Whitney tests, Control vs. cKO: AR, adjusted p = 0.7190; AU, adjusted p = 0. 0.7190; IN, adjusted p = 0. 0.7190. All data presented as mean ± SEM.

To further probe knockout effect on motivation, mice were tested on PR3 and PR7 reinforcement schedules, where each subsequent pellet required additional “active” lever presses. Overall pellet acquisition during PR sessions was comparable between genotypes (**Fig. 2b**). Yet, female *Grin1* cKO mice earned more pellets than female controls in PR3 (**Fig. 2c**), while males again showed no difference (**Fig. 2d**).

To determine whether this effect was driven by indiscriminate or intentional lever pressing, we quantified “active rewarded” (AR) presses, “active unrewarded” (AU) presses during time-outs, and “inactive” (IN) presses across the first 10 FR5 sessions. Total presses in each category were similar between genotypes (**Fig. 2e**). However, female *Grin1* cKO mice generated more AR presses than female controls (**Fig. 2f**), while male performance remained unchanged across genotypes (**Fig. 2g**). Both sexes, regardless of genotype, pressed the AR lever far more than AU or IN levers (**Fig. 2e–g**). Together, these results indicate that female *Grin1* cKO mice exhibit enhanced operant responding, likely reflecting increased hunger and/or reward motivation, rather than generalized hyperactivity or indiscriminate lever pressing.

### Impaired NMDAR-mediated glutamatergic input to ALDH1A1^+^ midbrain DANs does not reduce willingness to work for a high-effort reward

To further assess motivation and effort-based decision-making, we implemented a cost–benefit operant task during select FR5 and FR10 sessions (**Fig. 2a**). In this paradigm, mice were given two options: (1) exert greater effort to obtain high-value chocolate pellets, or (2) consume freely available but lower-value standard chow [21, 36]. Both *Grin1* cKO and control mice, regardless of sex, consumed only minimal amounts of freely available chow, even under different session conditions such as pellet limits (**Supplementary Fig. 4a-s**). This indicates that disrupting NMDAR-mediated glutamatergic input to ALDH1A1^+^ DANs did not impair motivation to work for the preferred reward. The consistently low chow consumption across FR5 and FR10 sessions suggests that all mice maintained a strong preference for and willingness to exert effort toward obtaining chocolate pellets, despite the availability of an easier, less desirable alternative.

### Impaired NMDAR-mediated glutamatergic input to ALDH1A1^+^ midbrain DANs is associated with sex-specific increases in food consumption and weight gain following food-restriction

Given the female-specific increases in pellet acquisition during entire operant learning, we next asked whether *Grin1* cKO mice display excessive or binge-like eating. Because operant tasks involved daily food restriction, we hypothesized that repeated restriction might have induced an abnormal drive to eat beyond what is required to restore metabolic balance. To test this, we examined free feeding behavior and body weight changes following food restriction.

After two consecutive nights of restriction, mice were housed individually in cages equipped with automated feeders [37] dispensing grain-based pellets nutritionally matched to standard chow (**Fig. 3a**). Feeders provided continuous access, one pellet at a time, enabling precise quantification of consumption. After six days, feeders were removed, and standard chow was restored. Body weights and pellet consumption were assessed separately in females and males.

**Fig. 3.**
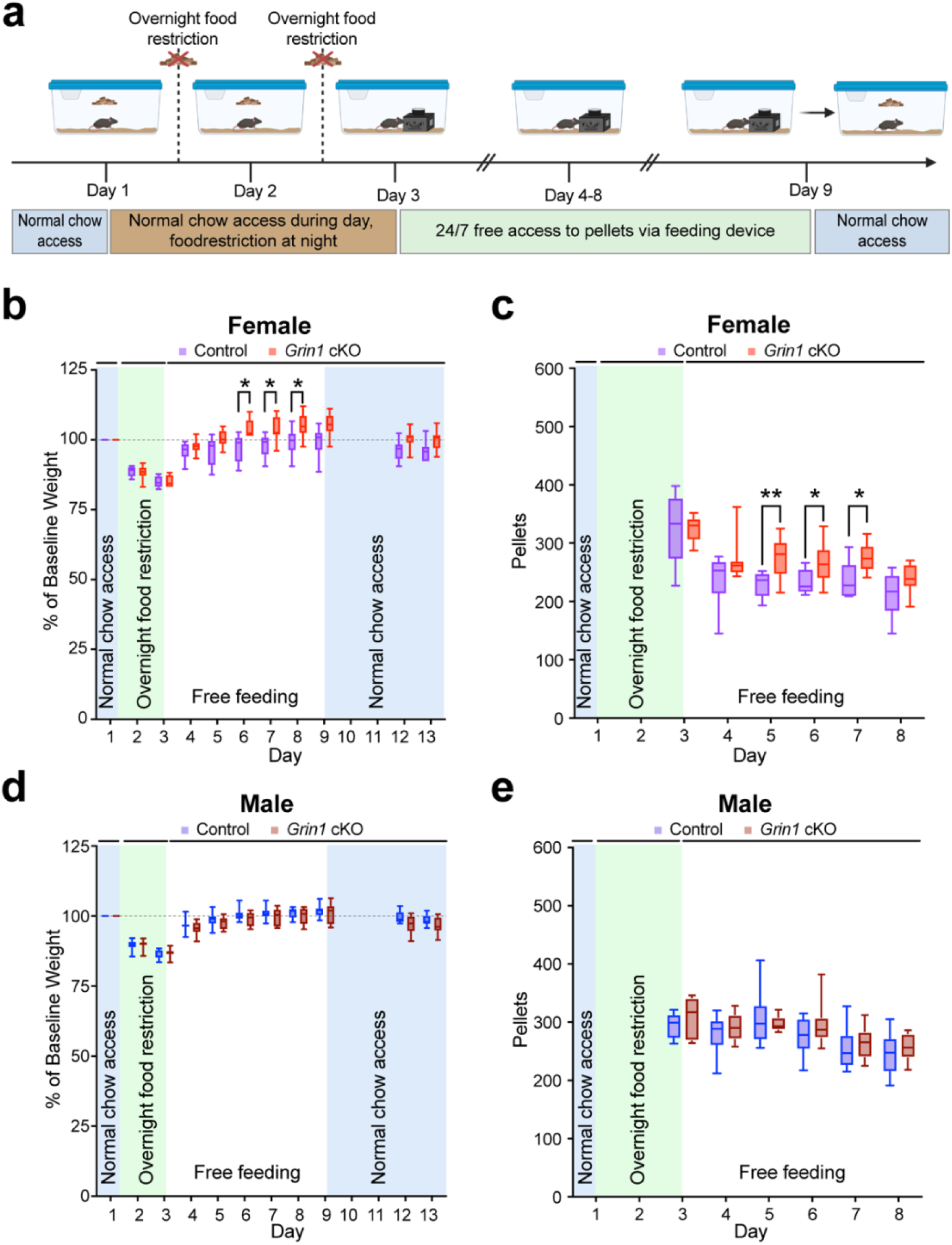
Impaired NMDAR-mediated glutamatergic input to ALDH1A1^+^ DANs causes transient overeating and weight gain in female *Grin1* cKO mice following food restriction. a, Timeline of overnight food restriction and free feeding using automated feeders, created by BioRender. **b**, Average percent baseline body weight of female *Grin1* cKO (n = 10) and control mice (n = 6). Mixed-effects analysis, genotype: F(1,14) = 7.542, *p = 0.0158; Tukey’s multiple comparisons test, day 6, *p = 0.0204; day 7, *p = 0.0266; day 8, *p = 0.0443. |**c**, Average daily pellet consumption of female *Grin1* cKO and control mice across the experiment. Mixed-effects analysis, genotype: F(1,14) = 4.622, *p = 0.0495; Tukey’s multiple comparisons test, day 5, **p = 0.0078; day 6, *p = 0.0353; day 7, p = 0.0332. **d**, Average percent baseline body weight of male *Grin1* cKO (n = 10) and control (n = 10) mice. Mixed-effects analysis, genotype: F(1,18) = 1.896, p = 0.1854. **e**, Average daily pellet consumption of male *Grin1* cKO and control mice. Mixed-effects analysis, genotype: F(1,18) = 1.720, p = 0.2062. All data are shown as box-and-whisker plots (min, first quartile, median, third quartile, max).

In females, both genotypes lost weight after restriction. Control females regained baseline weight within days of free feeding (**Fig. 3b**). In contrast, female *Grin1* cKO mice not only regained lost weight more rapidly but exceeded baseline values, showing a transient overshoot (**Fig. 3b**). Two weeks after the automated feeders had been removed, the average percent baseline body weights of female *Grin1* cKO became similar to controls (control: 98.00%, *Grin1* cKO: 100.8%; unpaired t-test with Welch correction, p = 0.1974). This divergence in weight gain in the days following restriction was accompanied by significantly greater pellet consumption in female *Grin1* cKO mice, both overall and particularly on Days 5–7 of free feeding (**Fig. 3c**).

In males, *Grin1* cKO and controls showed similar body weight decreases during restriction, followed by recovery to baseline within a few days of free feeding before stabilizing (**Fig. 3d**). Body weight remained comparable two weeks later (control: 97.40%, *Grin1* cKO: 94.76%; unpaired t-test with Welch correction, p = 0.1448). Daily pellet consumption also did not differ between male groups (**Fig. 3e**). Together, these findings reveal a female-specific vulnerability, in which inhibiting NMDAR-mediated glutamatergic input to ALDH1A1^+^ neurons promotes transient overeating and weight gain following food restriction.

### Regionally specific viral *Grin1* knockdown reduces GluN1 in ALDH1A1^+^ SNc and VTA DANs

We next examined whether NMDAR-mediated glutamatergic input to ALDH1A1^+^ DANs in the SNc or VTA contributes to the altered feeding observed in female *Grin1* cKO mice. To achieve region-specific *Grin1* knockdown (KD), *Aldh1a1*^+/P2A-CreERT2^ mice received bilateral stereotaxic injections of Cre-dependent AAVs encoding SaCas9 and sg*Grin1* [38] into either the SNc (SNc *Grin1* KD) or VTA (VTA *Grin1* KD) (**Fig. 4a**). Control groups received Cre-dependent SaCas9 without guide RNA (SaCas9-empty) into the same regions (SNc control, VTA control). The *Grin1* KD construct carried an HA tag, allowing identification of transduced cells (**Supplementary Fig. 5a**). Viral transduction was significantly enriched in ALDH1A1^+^ DANs within the targeted region compared to the non-targeted region in both SNc and VTA *Grin1* KD mice. In SNc *Grin1* KD brains, 67.8% of all HA^+^ neurons localized to the SNc, whereas in VTA *Grin1* KD brains, 67.5% of all HA^+^ neurons localized to the VTA (**Supplementary Fig. 5b**). Immunohistochemistry confirmed regionally specific *Grin1* KD with GluN1 protein levels reduced in ALDH1A1^+^ neurons, but not ALDH1A1^-^ neurons, within the targeted region (**Fig. 4b**). In contrast, SaCas9-empty controls showed no GluN1 reduction in ALDH1A1^+^ neurons (**Fig. 4b**).

**Fig. 4.**
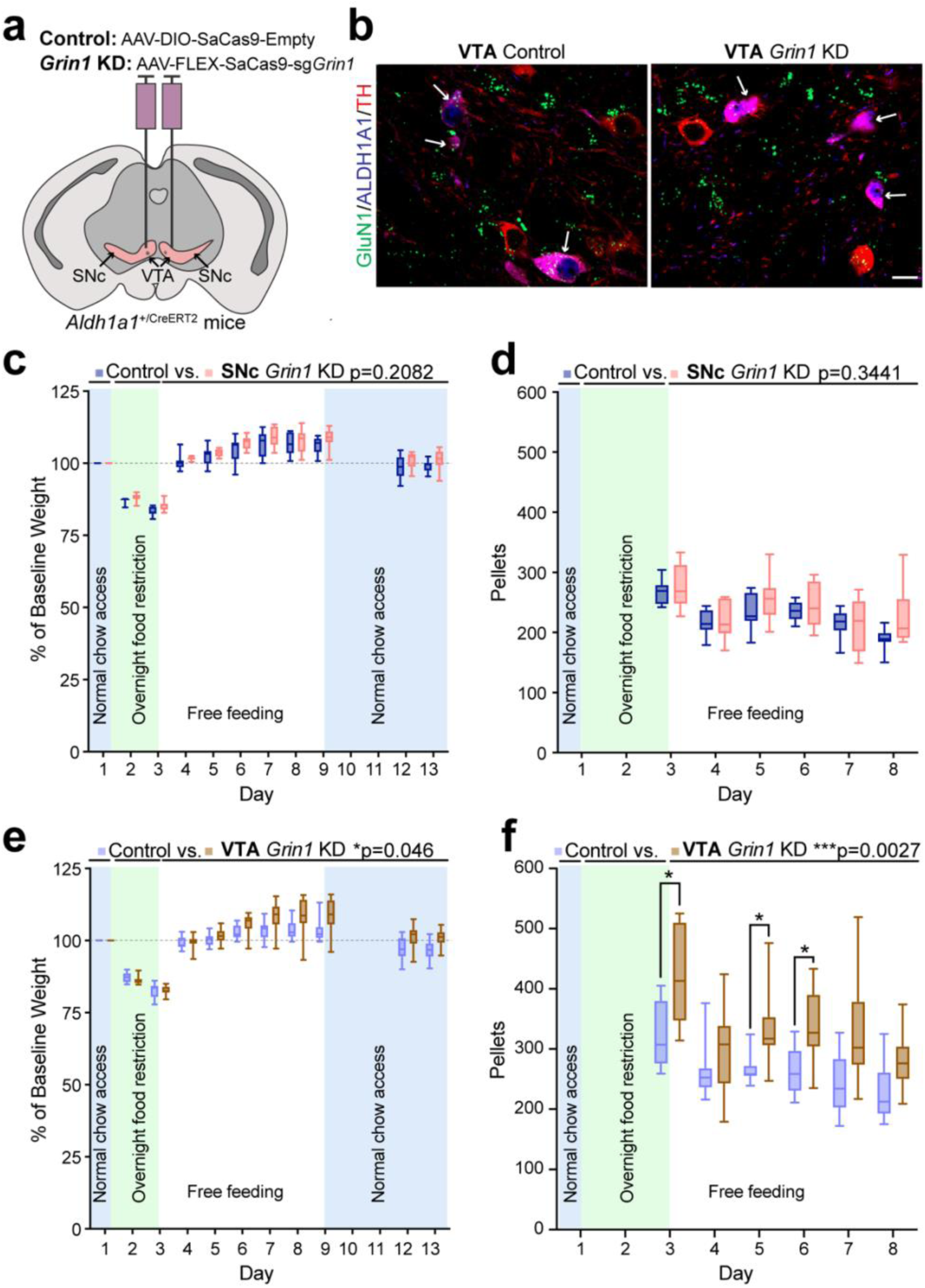
Regionally specific knockdown of *Grin1* in ALDH1A1^+^ VTA DANs reproduces post-restriction increases in food intake and body weight in females. a, Brain schematic showing bilateral AAV injection sites in the VTA of *Aldh1a1*^+/P2A-CreERT2^ mice. **b**, Representative confocal images (63×, single z-plane) of TH (red), ALDH1A1 (blue), and GluN1 (green) co-staining in coronal VTA sections from *Aldh1a1*^+/P2A-CreERT2^ mice injected with AAV9-FLEX-SaCas9-sg*Grin1* (VTA *Grin1* KD) or AAV9-DIO-SaCas9-empty (VTA control). White arrows indicate ALDH1A1^+^ DANs. Scale bar, 20 μm. **c**, Average percent baseline body weight of female SNc *Grin1* KD (n = 8) and control (n = 17) mice. Mixed-effect analysis, genotype: F(1,13) = 1.754, p = 0.2082. **d**, Average daily pellet consumption of female SNc *Grin1* KD (n = 8) and control mice (n = 7). Mixed-effect analysis, genotype: F(1,13) = 0.9640, p = 0.3441. **e,** Average percent baseline body weight of female VTA *Grin1* KD (n = 11) and control (n = 10) mice. Mixed-effect analysis, genotype: F(1, 19) = 4.565, *p = 0.0458. Multiple comparisons test, day 3, adjusted p > 0.9999; day 4, adjusted p > 0.9999; day 5, adjusted p > 0.9999; day 6, adjusted p = 0.3699; day 7, adjusted *p = 0.0208; day 8, adjusted *p = 0.0291; day 9, adjusted *p = 0.0122; day 12, adjusted p = 0.1093; day 13, adjusted p = 0.0825. **f,** Average daily pellet consumption of female VTA *Grin1* KD and control mice. Mixed-effect analysis, genotype: F(1, 19) = 11.87, ***p = 0.0027. Multiple comparisons test, day 3, adjusted **p = 0.0040; day 4, adjusted p = 0.1617; day 5, adjusted **p = 0.0053; day 6, adjusted **p = 0.0021; day 7, adjusted *p = 0.0211; day 8, adjusted *p = 0.0147. All data shown as box-and-whisker plots (min, first quartile, median, third quartile, max).

### Impaired NMDAR-mediated glutamatergic input to ALDH1A1^+^ VTA DANs reproduces post-restriction increases in food consumption and weight gain in female mice

We next used the SNc and VTA *Grin1* KD models to determine whether disruption of NMDAR-mediated input to ALDH1A1^+^ neurons in either region contributes to the abnormal feeding and weight gain observed in female *Grin1* cKO mice. Locomotor behavior was first assessed in the open field assay. Neither SNc nor VTA *Grin1* KD females differed from controls in distance traveled, speed, or center versus border time (**Supplementary Fig. 6**), confirming no locomotor deficits.

Following food restriction, SNc *Grin1* KD and control females showed similar decreases in body weight, recovery to baseline upon refeeding, and stable long-term weights (**Fig. 4c**). Two weeks after feeder removal, their average percent baseline body weights remained comparable (SNc control: 103.2%, SNc *Grin1* KD: 103.0%; unpaired t-test with Welch correction, p = 0.8774). Pellet intake during free feeding was likewise indistinguishable (**Fig. 4d**).

In contrast, VTA *Grin1* KD females displayed marked alterations. After food restriction, they regained weight rapidly and overshot baseline, recapitulating the phenotype of *Grin1* cKO females (**Fig. 4e**). This overshoot was transient with weights normalizing within two weeks after feeder removal (VTA control: 100.1%, VTA *Grin1* KD: 103.1%; unpaired t-test with Welch correction, p = 0.0969). During free feeding, VTA *Grin1* KD females also consumed significantly more pellets than controls (**Fig. 4f**). Together, these findings demonstrate that impaired NMDAR-mediated glutamatergic input to ALDH1A1^+^ VTA, but not SNc, DANs is sufficient to induce transient overeating and excessive weight gain in female mice following food restriction.

### *Grin1* cKO versus control comparisons reveal major gene expression differences in female food-restricted mice

Hunger and satiety signals influence dopaminergic reward circuits by modulating neuronal activity and reward sensitivity [39]. To investigate how disrupted NMDAR-mediated glutamatergic input to ALDH1A1^+^ DANs interacts with metabolic state, we performed whole brain transcriptomic profiling across genotype, sex, and feeding condition. Three feeding groups were examined: (1) no restriction, (2) post-restriction (two nights of food restriction), and (3) refeeding (two nights of restriction followed by 72 h of *ad libitum* access via automated feeder) (**Fig. 5a**).

**Fig. 5.**
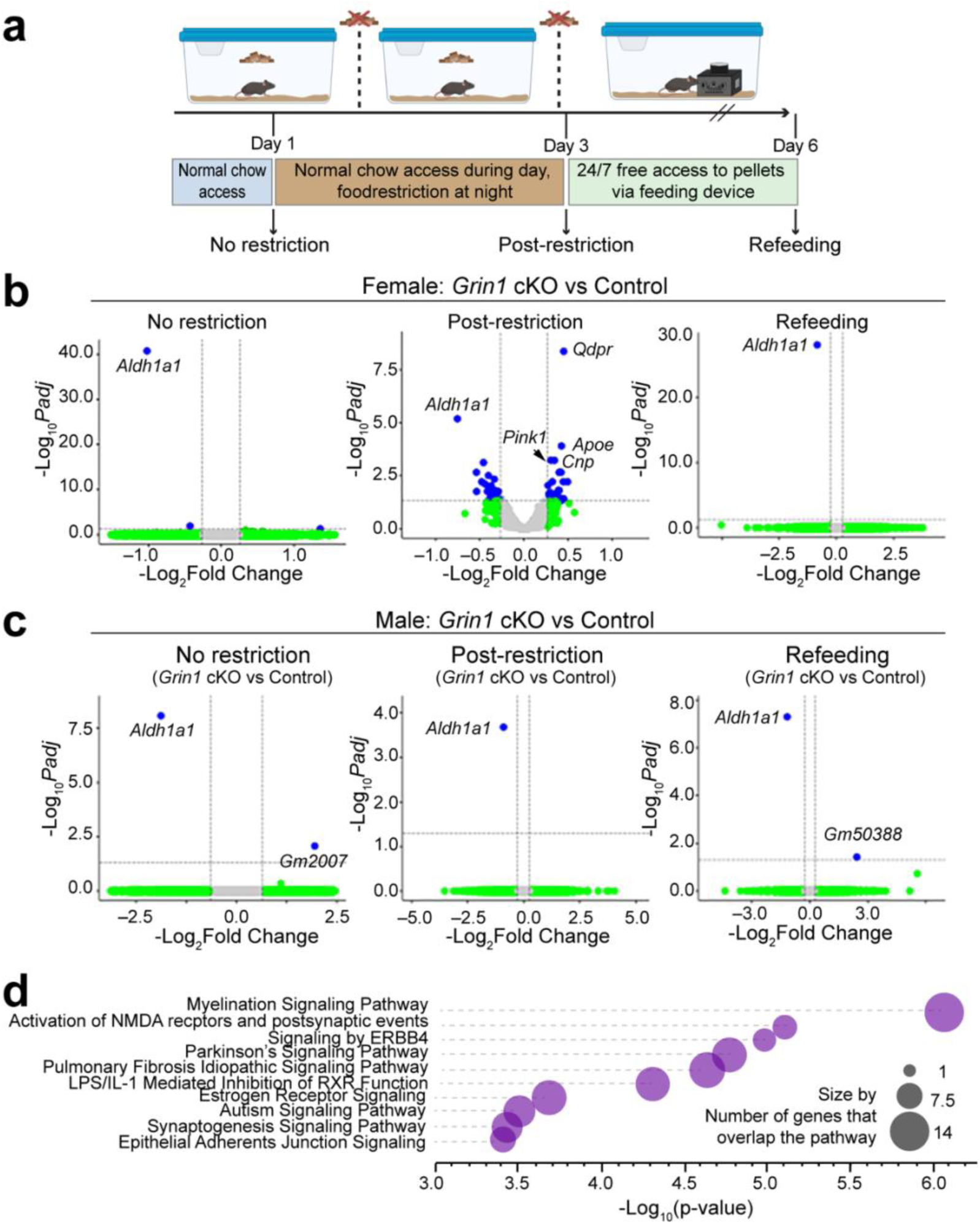
*Grin1* cKO versus control comparisons reveal broad transcriptional differences in female food-restricted mice. **a**, Experimental timeline showing food restriction, refeeding, and mRNA-seq collection timepoints. Male and female *Grin1* cKO and control mice were sampled at three conditions: no restriction, post-restriction, and refeeding. Created with BioRender. **b–c**, Enhanced volcano plots of differentially expressed genes (DEGs) between *Grin1* cKO and control mice for each feeding condition, separated by sex. Gray points: non-significant genes; green: fold change > 1.2 but adjusted p > 0.05; blue: significant DEGs (adjusted p < 0.05, fold change > 1.2). Genes to the right are upregulated in *Grin1* cKO versus control; those to the left are downregulated. padj, adjusted p-value. **d**, Pathway enrichment analysis of DEGs in female *Grin1* cKO versus control post-restriction mice. Bubble plot shows the top ten enriched pathways. Bubble size reflects number of overlapping DEGs.

Strikingly, transcriptional differences emerged almost exclusively between female *Grin1* cKO and control mice after food restriction (**Fig. 5b–c, Supplementary Table 1**). This indicates that gene expression changes were not a simple consequence of *Grin1* loss but instead reflected the combined influence of sex, metabolic stress, and disrupted NMDAR signaling. Notably, *Aldh1a1* expression was markedly reduced in both male and female Grin1 cKO mice under all three experimental conditions (**Fig. 5b, 5c**). This reduction likely results from impaired *Aldh1a1* transcription due to insertion of the *P2A-CreERT2* cassette into the 3’ untranslated region of the *Aldh1a1* gene locus in *Aldh1a1*^+/P2A-CreERT2^ mice [14].

Pathway analysis revealed altered expression of genes linked to NMDA receptor activation, postsynaptic signaling events, and Parkinson’s disease–associated pathways (**Fig. 5d**). These results highlight a context-specific and female-biased transcriptional response, suggesting enhanced vulnerability or adaptive plasticity of ALDH1A1^+^ DANs under energetic stress.

### Female versus male comparisons reveal major genotype-dependent transcriptional differences in *Grin1* cKO food-restricted mice

To determine whether disrupted NMDAR-mediated glutamatergic input produces sex-dependent transcriptional effects, we compared female and male mice within each genotype across three feeding conditions. In control mice, there were no differentially expressed genes (DEGs) between females and males in either the post-restriction or refeeding groups, and only a few DEGs in the no restriction group (**Fig. 6a**). By contrast, *Grin1* cKO mice exhibited robust sex-specific transcriptional differences (**Fig. 6b**). In the post-restriction group, female versus male *Grin1* cKO mice exhibited 124 upregulated and 92 downregulated genes (**Supplementary Table 2**). Differences were even more pronounced in the refeeding group with 388 upregulated and 82 downregulated genes (**Supplementary Table 3**). Notably, 82.6% of DEGs in the refeeding comparison were upregulated, compared with 57.4% in the post-restriction comparison and 48.5% in the female *Grin1* cKO versus control post-restriction comparison.

**Fig. 6.**
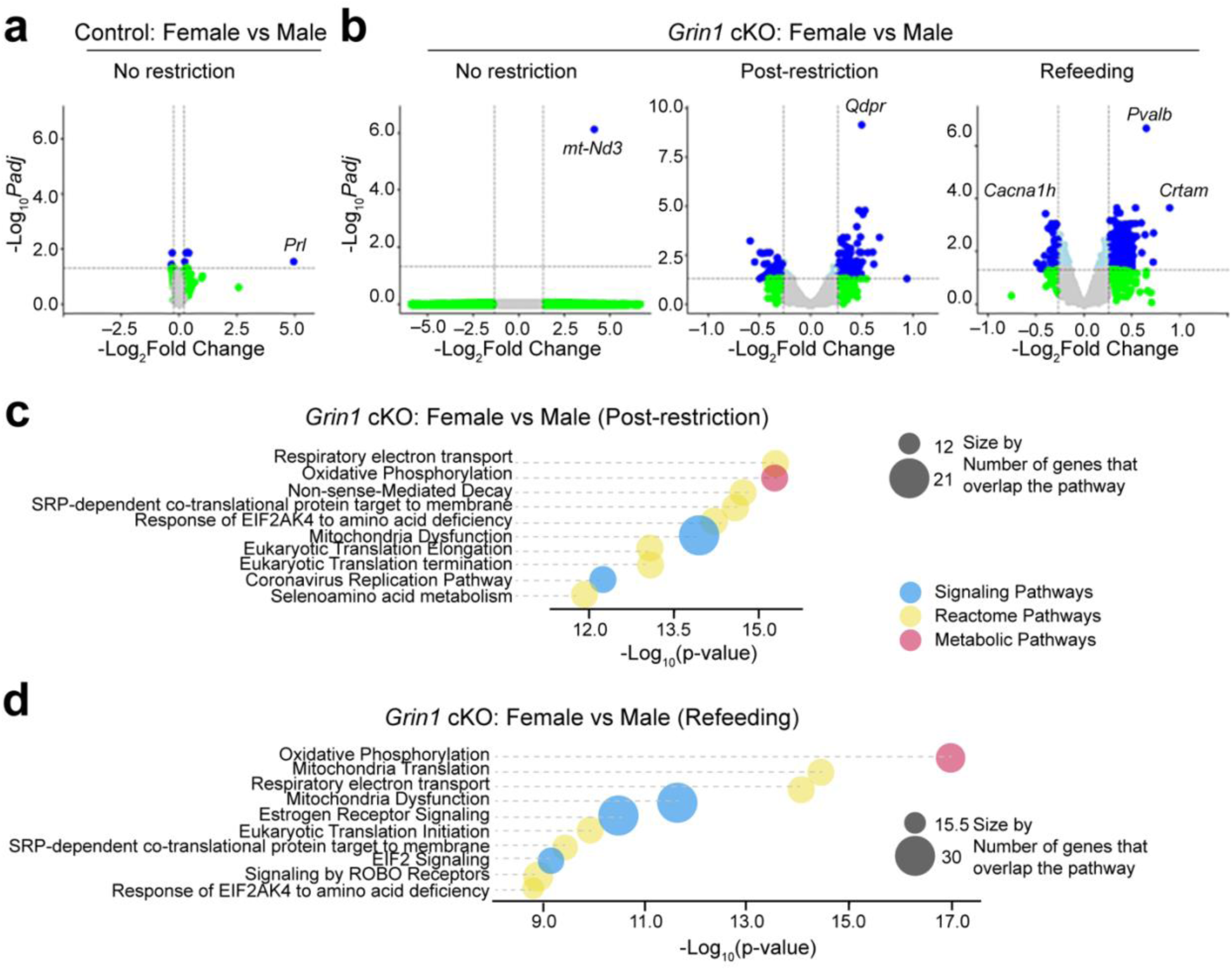
Female versus male comparisons reveal sex-dependent transcriptional differences in *Grin1* cKO food-restricted mice. **a-b**, Enhanced volcano plots of DEGs between female and male mice across feeding conditions for *Grin1* cKO and control groups. Colors and fold-change criteria as in Fig. 5. **c–d**, Pathway enrichment analysis of DEGs between female and male *Grin1* cKO mice under post-restriction (b) and refeeding (c) conditions. Bubble plots display the ten most significantly enriched pathways. Bubble size represents the number of overlapping DEGs; color denotes pathway category (blue, Signaling; yellow, Reactome; red, Metabolic).

Pathway enrichment analysis revealed that many DEGs between female and male *Grin1* cKO mice mapped to energy and metabolism-related pathways. In the post-restriction group, the top five enriched pathways were: (1) Respiratory Electron Transport, (2) Oxidative Phosphorylation, (3) Nonsense-Mediated Decay, (4) SRP-dependent Co-translational Protein Targeting to Membrane, and (5) EIF2AK4 (GCN2) Response to Amino Acid Deficiency (**Fig. 6c**). In the refeeding group, the top five were: (1) Oxidative Phosphorylation, (2) Mitochondrial Translation, (3) Respiratory Electron Transport, (4) Mitochondrial Dysfunction, and (5) Estrogen Receptor Signaling (**Fig. 6d**). Importantly, Respiratory Electron Transport and Oxidative Phosphorylation consistently ranked among the top three pathways in both conditions, highlighting a sex-dependent effect of *Grin1* loss on mitochondrial and metabolic processes.

Taken together, these findings suggest that female *Grin1* cKO mice undergo unique transcriptional adaptations in metabolic and mitochondrial pathways under energetic stress, which may contribute to their enhanced food consumption and weight gain following food restriction.

## Discussion

PD is increasingly recognized as a multisystem disorder with both motor and non-motor components, including compulsive and dysregulated eating behaviors that are more prevalent in women [2, 4]. To understand the neural mechanisms linking dopaminergic dysfunction to such metabolic phenotypes, we used sophisticated genetic and behavioral approaches to selectively disrupt NMDAR-mediated glutamatergic signaling in ALDH1A1^+^ DANs and found that female, but not male, *Grin1* cKO mice exhibit excessive feeding with transient weight gain following food restriction. Region-specific *Grin1* knockdown identified the VTA, rather than the SNc, as the critical site underlying this phenotype. Transcriptomic profiling further uncovered sex-dependent gene expression changes related to mitochondrial function, energy metabolism, and synaptic signaling, defining a sex-specific mechanism by which excitatory regulation of ALDH1A1^+^ VTA DANs modulates feeding behavior in PD.

Consistent with prior work showing that many dopamine-associated behaviors remain intact despite disruption of NMDAR-mediated burst firing in midbrain DANs [19, 20], we found that selective loss of NMDAR input to ALDH1A1^+^ DANs did not impair locomotor activity or rotarod motor skill learning. This differs from studies where ablation or inhibition of ALDH1A1^+^ SNc DANs impaired motor performance [14, 40], suggesting that loss of NMDAR-mediated glutamatergic input is less disruptive than complete neuronal loss. Compensatory excitatory or inhibitory inputs may preserve function, or homeostatic mechanisms may adapt to reduced glutamatergic drive. Thus, the *Grin1* cKO model, which reduces excitatory regulation without ablation or inhibition, may be valuable for probing early dopaminergic dysfunction in PD, when neurons remain intact but signaling is altered.

Unlike prior studies showing that combined disruption of NMDAR and AMPAR inputs impairs motivated behavior [21], we observed no deficits in operant tasks requiring effort-based responding in *Grin1* cKO mice. Both males and females maintained normal motivation to work for preferred rewards under increasingly demanding operant schedules. Instead, female *Grin1* cKO mice showed enhanced food acquisition compared with controls, particularly during early FR1 training when minimal effort was required, suggesting a sex-specific increase in feeding drive. The absence of elevated inactive lever pressing further indicates that this effect did not result from indiscriminate responding.

Female-specific effects extended to the free-feeding after food restriction, where *Grin1* cKO females consumed more food and displayed transient weight gain above baseline. Region-specific viral *Grin1* knockdown demonstrated that disruption in the VTA, but not the SNc, was sufficient to reproduce this phenotype. This finding is consistent with the established role of VTA dopaminergic circuits in motivation and reward-related behaviors, including feeding, where nucleus accumbens dopamine release promotes palatable food consumption [41–43]. These results suggest that impaired glutamatergic regulation of ALDH1A1^+^ VTA DANs amplifies both homeostatic and hedonic drive to eat following energy depletion.

Transcriptomic analyses provided a mechanistic link between disrupted excitatory input and sex-specific behavioral outcomes. Gene expression changes were minimal between male controls and *Grin1* cKO mice but were pronounced between female controls and *Grin1* cKO mice following food restriction. Many DEGs mapped to NMDAR signaling, postsynaptic plasticity, and PD–associated pathways, consistent with selective disruption of NMDAR signaling in PD-vulnerable ALDH1A1^+^ DANs. Comparisons between female and male *Grin1* cKO mice identified hundreds of DEGs enriched for oxidative phosphorylation, respiratory electron transport, and mitochondrial dysfunction. These pathways represent well-established early sites of PD vulnerability, where impaired bioenergetics and altered synaptic integration contribute to dopaminergic degeneration. The additional enrichment of estrogen receptor signaling in refed females suggests hormonal modulation of these adaptations.

Together, these findings demonstrate that NMDAR-mediated glutamatergic regulation of ALDH1A1^+^ VTA DANs plays a sex-specific role in integrating metabolic and reward signals. Female *Grin1* cKO mice not only displayed excessive feeding and transient weight gain but also exhibited transcriptional signatures consistent with PD-relevant vulnerabilities, including altered mitochondrial pathways and synaptic transmission. These convergent results highlight how sex and energetic state interact to shape DAN function and suggest that disrupted glutamatergic regulation of ALDH1A1^+^ DANs may contribute to compulsive eating disorders and sex differences in PD.

## Methods

### Mouse work

All animal procedures were approved by the Institutional Animal Care and Use Committee (IACUC) of the Intramural Research Program at the National Institute on Aging (NIA), National Institutes of Health (NIH), and conducted in accordance with NIH guidelines. Mice were maintained in a 12-hour light/12-hour dark cycle in an NIH animal facility with *ad libitum* access to water. All mice were group housed except during food restriction protocols. Aside from overnight food restriction during operant lever press and free feeding experiments, mice had continuous access to standard chow.

All behavioral testing, including open field, rotarod, and operant lever press tasks, was conducted during the light phase. The free feeding assay was performed continuously across both light and dark cycles for approximately ten days. The sex distribution of mice for each experiment is reported in the corresponding figure legends.

### *Grin1* cKO mice

An inducible Cre recombinase (CreERT2) knock-in mouse line expressing CreERT2 in ALDH1A1^+^ midbrain DANs (*Aldh1a1*^+/P2A-CreERT2^) was generated as previously described [14]. That line was crossed with floxed NR1 (*Grin1*^fl/fl^) mice (The Jackson Laboratory, strain #005246), which contain *loxP* sites flanking the transmembrane domain and C-terminal region of the *Grin1* gene. The resulting *Aldh1a1*^+/P2A-CreERT2^::*Grin1*^fl/fl^ conditional knockout (*Grin1* cKO) mice enable selective deletion of the NR1/GluN1 subunit of NMDA receptors in ALDH1A1^+^ midbrain DANs. *Grin1*^fl/fl^ and *Grin1*^fl/+^ mice were used as controls.

Male and female *Grin1* cKO and control mice were tested in open field, rotarod, operant lever press, and food restriction/free feeding behavioral assays. For open field and rotarod experiments, two age cohorts were tested: one at ∼3–4 months and another at 15–27 months. For operant and feeding experiments, mice were approximately 1 year old at the time of testing. For mRNA sequencing, mice were 6–8 months old at the time of brain collection.

All mice received 4-hydroxytamoxifen (4-OHT) injections for five consecutive days prior to testing to induce CreERT2-mediated recombination.

### Viral CRISPR/SaCas9 *Grin1* KD mice

The same *Aldh1a1*^+/P2A-CreERT2^ knock-in mouse line that was used to generate the conditional knockout was also used to generate viral *Grin1* knockdown (KD) cohorts. Mice received bilateral stereotaxic injections of Cre-dependent AAVs expressing SaCas9 with a single guide RNA (sgRNA) targeting *Grin1* (AAV-FLEX-SaCas9-U6-sg*Grin1*; Addgene #124852) [38] or a control virus expressing SaCas9 without sgRNA (AAV-DIO-SaCas9-U6-empty). AAVs were injected into either the SNc (SNc *Grin1* KD and control) or VTA (VTA *Grin1* KD and control) of female mice to selectively disrupt NMDAR-mediated glutamatergic input to ALDH1A1^+^ DANs within each region.

Female *Grin1* KD and control mice were tested in open field and food restriction/free feeding behavioral paradigms. Following viral injections, all mice received 4-OHT for five consecutive days to activate CreERT2. Behavioral testing was conducted when mice were approximately 6–7 months old.

### Stereotactic injection

Stereotactic survival surgery was performed under aseptic conditions in accordance with the Animal Research Advisory Committee (ARAC) guidelines for Rodent Survival Surgery. Adult mice were anesthetized with isoflurane and secured in a stereotaxic frame (Stoelting). Oxygen and isoflurane levels were continuously monitored throughout the procedure. Ophthalmic ointment was applied to protect the eyes, and hair over the scalp was shaved or trimmed. The surgical site was disinfected twice with iodine followed by 30% ethanol prior to incision.

A midline scalp incision was made to expose the skull, ensuring both bregma and lambda were visible. The head was leveled such that bregma and lambda were within 0.05 mm of each other in both medial–lateral and dorsal–ventral planes. Small craniotomies were drilled at the appropriate coordinates for viral injection. For substantia nigra pars compacta (SNc) targeting, coordinates relative to bregma were: anterior/posterior (A/P) −3.1 mm, medial/lateral (M/L) ±1.5 mm, and dorsal/ventral (D/V) −3.9 mm from the dura. For ventral tegmental area (VTA) targeting, coordinates were: A/P −3.4 mm, M/L ±0.4 mm, and D/V −4.2 mm from the dura.

For each injection, 250–500 nL of the desired AAV (titer 1.29 × 10^13^–2.75 × 10^13^ GC/mL) was loaded into a 2.0 µL syringe (Hamilton NEUROS model 53495) and delivered at 75 nL/min using a motorized stereotaxic injector (Stoelting). After completion of the injection, the needle was left in place for 5 minutes before slow withdrawal. All injections were performed bilaterally.

Following injection, topical antibiotics were applied, and the incision was closed using tissue adhesive (3M™ Vetbond™ 1469SB) and/or silk sutures (DemeSILK™, 5-0, DT-719-1, DemeTECH). Additional antibiotic ointment was applied to the wound. Mice received subcutaneous injections of extended-release meloxicam (6 mg/kg) and 1 mL sterile saline for analgesia and hydration. Animals were returned to their home cages with a heat pack and monitored until full recovery from anesthesia. Postoperative monitoring by veterinary staff continued daily to assess for pain or distress.

### Cre induction with tamoxifen

(Z)-4-Hydroxytamoxifen (4-OHT) (Sigma-Aldrich, H7904) was dissolved in ethanol (10 mg/1200 µL). Then, two times the volume of corn oil (C8267, Sigma-Aldrich) was added until completely dissolved. The solution was aliquoted into microcentrifuge tubes and ethanol was vaporized using a vacuum centrifuge for 20-30 minutes. Additional corn oil was added to make a 1 mg/ml solution. The solution was then filtered using a 50 mL tube top vacuum filter system (430320, Corning), aliquoted, and stored at −20°C. Mice were injected intraperitoneally for 5 consecutive days with the 1 mg/ml 4-OHT solution at a dose of 10 mg/kg body weight (injection volume of 10 µL/g). The cKO and control mice were at least 2 months of age before injections and CRISPR viral KD mice were injected at least 1 week after surgery. Mice were given at least 4 weeks post-4-OHT injection before any testing or tissue collection was performed to ensure sufficient 4-OHT-induced Cre-loxP recombination.

### Open field spontaneous locomotion test

Mice were acclimated for at least an hour to the behavior room prior to testing. The open field test occurred in the dark with a diffuse source of light. Prior to adding mice to the open field arenas, videos of the empty arenas were obtained for background calibration during video analysis. The mice were then placed in a 43.815 cm × 43.815 cm × 39.37 cm (length × width × height) white arena, and their voluntary activity was recorded (WinTV v7) for 5 minutes with overhead video tracking. Before and after each arena was used for testing, the arena was cleaned thoroughly with 70% alcohol. Videos were then analyzed using TopScan software (version 3), which tracked each mouse’s nose to determine its position and velocity throughout the experiment. For additional analysis, the arena was divided into a center and border portion based on previously utilized ratios [14]. The border included the area 5.84 cm in from each edge while the center included the remaining area. Arenas were individually calibrated for each trial. Data on each mouse’s position, distance traveled, and velocity at each 1 second bin throughout the session was exported and used to determine the distance traveled in the border and center of the arena and in total, the average velocity in the border and center of the arena, and the time spent in the border and center of the arena. In total, 3 mice failed to spend any time in the center of the open field arena and were therefore excluded from analysis (1 male control in the 3-4 month-old cohort, 1 male control in the 15-27-month-old cohort, and 1 male cKO in the 15-27-month-old cohort).

### Rotarod motor learning task

Performance on a rotarod apparatus (Model LE8205, Panlab/Harvard Apparatus) was used to assess motor skill learning. Mice are placed on a grooved rod (30 mm diameter) that accelerates from 0 to 40 in 5 minutes. Performance was measured as the latency for each mouse to fall off the rotating rod. The trial ended when the mouse fell off the rotarod or after 300 seconds, whichever came first. Each mouse completed 10 trials a day for 6 consecutive days. Between the end of one trial and the start of the next trial, mice were given an approximately 2-minute break. Throughout the test, any mouse urine and feces were removed to keep rotarod surfaces clean and dry.

### Lever presses operant conditioning task

Fixed ratio (FR) and progressive ratio (PR) reinforcement schedules were used to assess operant learning and motivation to work for a reward under progressively demanding conditions. Each mouse’s free feeding body weight was recorded prior to food restriction and monitored daily throughout the experiment. During the week preceding testing, mice were food restricted for five consecutive nights, and subsequently overnight prior to each operant session to ensure task motivation. This regimen maintained mice at ∼90% of their free-feeding body weight throughout the experiment. Following each session, mice were provided ad libitum access to standard chow until the next overnight restriction period.

Behavioral testing was conducted in sound-attenuated operant chambers with grid floors (ENV-307W, Med Associates) controlled by MED-PC IV software. Each chamber contained two retractable levers (ENV-312-2W, Med Associates) positioned on either side of a pellet trough connected to a pellet dispenser. A house light and ventilation fan provided illumination and white noise throughout each session. For each mouse, one lever was designated as the “active” lever, while the other served as the “inactive” lever. Pressing the active lever delivered a 20 mg chocolate-flavored pellet (F05301, Bio-Serv), followed by a 5 s time-out during which additional active lever presses did not yield rewards. Presses on the “inactive” lever had no programmed consequence.

### Pre-training and Fixed Ratio 1

Mice first completed four days of 45-minute pre-training sessions. During these sessions, both levers were extended, but lever presses had no programmed consequence. Chocolate pellets were delivered at random variable intervals (mean 45 s; range 4–132 s).

Mice then completed ten daily sessions on a fixed ratio 1 (FR1) schedule, in which each “active” lever press (outside the 5 s time-out period) resulted in delivery of a pellet reward. Each session lasted 45 minutes or until 50 pellets were earned, whichever occurred first. Only mice that earned ≥40 pellets in at least one session were included in the analysis. Most mice reached the 50-pellet maximum within several days. Out of 36 total mice, two (1 female experimental and 1 male control) failed to acquire the lever-pressing behavior and were excluded. The number of pellets earned per session was recorded.

### Choice/No Choice task (Fixed Ratio 5 and Fixed Ratio 10)

To evaluate effort-based decision-making, a Choice/No Choice paradigm adapted from previous studies was used [21, 36]. Mice first underwent three days of pre-training. On pre-training Day 1, mice were tested on a fixed ratio 5 (FR5) schedule, where every five “active” lever presses resulted in a chocolate pellet reward. A pre-weighed cup of freely available standard chow was also placed in the chamber, allowing mice to choose between working for pellets or consuming chow (“FR5 Choice” session). On pre-training Day 2, mice completed an FR5 session without freely available chow (“FR5 No Choice”). On pre-training Day 3, mice were placed in the chamber with only freely available chow; lever presses had no consequence. All pre-training trials lasted 45 minutes or until 50 pellets were earned, whichever came first.

The following 20 sessions alternated between Choice and No Choice conditions. During the first ten sessions, mice completed FR5 Choice sessions on Days 1, 3, 5, 6, 8, and 10, and FR5 No Choice sessions on Days 2, 4, 7, and 9. Each session lasted 45 minutes or until 50 pellets were earned, whichever came first. To avoid ceiling effects, the 50-pellet limit was removed for the next five sessions, which each lasted 45 minutes (Choice: Days 11, 13, and 15; No Choice: Days 12 and 14).

Next, mice completed five days of FR10 sessions, where every ten “active” lever presses resulted in a reward. FR10 Choice sessions were conducted on Days 1, 3, and 5, and FR10 No Choice sessions on Days 2 and 4. All FR10 sessions lasted 45 minutes with no pellet limit. For Choice sessions, the total weight of consumed food (pellets + chow) and the percentage of total intake from chocolate pellets were calculated.

### Progressive Ratio 3 and Progressive Ratio 7

After completing FR5 and FR10 sessions, mice were tested on progressive ratio (PR) schedules. Mice first completed five consecutive days on a PR3 schedule, in which the number of lever presses required to earn each subsequent pellet increased by three (4, 7, 10, 13, etc.). For the first three days, sessions lasted 45 minutes or ended after 5 minutes without a lever press, whichever came first. For the final two days, session duration was extended to 90 minutes with the same inactivity cutoff.

Mice then completed ten days on a PR7 schedule, where each subsequent pellet required seven additional lever presses. For the first seven days, sessions lasted 90 minutes or ended after 5 minutes of inactivity, whichever came first. On Days 8 and 9, session length was extended to 240 minutes, and on Day 10, to 420 minutes. The total number of pellets earned per session was recorded.

### Free feeding with FED3 devices

Food consumption and weight changes following food restriction were measured to assess post-restriction feeding behavior. At the start of the experiment, all mice were weighed to establish baseline body weights and then singly housed with *ad libitum* access to standard rodent chow and water. Between Day 1 and Day 2, food was removed overnight. Mice were weighed the following morning (end of restriction), and chow was returned. Food was again removed overnight between Day 2 and Day 3, after which mice were weighed and automatic Feeding Experimentation Device version 3 units (FED3; Open Ephys) [37] were placed in each cage. Throughout the entire experiment, mice had *ad libitum* access to water in their cage.

FED3 devices were set to free feed mode and loaded with 20 mg grain-based pellets (Dustless Precision Pellets®, #F0163, Bio-Serv). In this mode, a single pellet is dispensed into the pellet trough; a new pellet is only dispensed once the previous one is removed. Mice had continuous access to pellets via the FED3 devices for six consecutive 24-hour periods (Days 3–9). Each day at approximately 12:00 PM, devices were checked for jams, refilled as needed, and mice were weighed. If a jam occurred within the preceding 24-hour period, pellet and weight data from that day were excluded. One female VTA control mouse was excluded entirely from analysis due to the development of health issues. Event timestamps (e.g., pellet retrievals) were automatically logged to SD cards in CSV format.

After the sixth 24-hour session, FED3 devices were removed. Mice remained singly housed for four additional days (Days 9–13) with *ad libitum* chow. Weights were recorded on Days 12 and 13, and again two weeks later after mice were returned to group housing with standard chow and water.

To control for baseline differences, body weights were expressed as percentages of each mouse’s initial baseline weight. In figures, the average percent baseline weight for each day reflects the body weight recorded that day, while average pellet counts represent pellets consumed during the 24-hour period beginning on that day and ending the following day.

### Immunohistochemistry

Mice were anesthetized with pentobarbital (50–90 mg/kg, i.p.) and transcardially perfused with pre-chilled phosphate-buffered saline (PBS), followed by 4% paraformaldehyde (PFA; 15714-S, Electron Microscopy Sciences), using a peristaltic pump (GP1000, Fisher Scientific) at ∼15 mL/min. Briefly, once mice were unresponsive to tail and toe pinches, the thoracic cavity was opened, the heart exposed, and a 23-gauge infusion needle inserted into the left ventricle. A small incision was made in the right atrium. PBS was perfused for ∼1.5 minutes until the liver became pale, followed by 4% PFA for ∼2.5 minutes, and a final PBS rinse for ∼30 seconds. Successful fixation was indicated by body twitching and tail flicking.

Brains were removed and post-fixed overnight in 4% PFA at 4°C, then cryoprotected in 30% sucrose in PBS for at least two days before sectioning. Coronal sections (40 μm) were cut on a cryostat (Leica CM3050 S, Leica Biosystems) and stored in PBS containing 0.06% sodium azide (Sigma-Aldrich). Sections were blocked for 1 hour at room temperature (or overnight at 4°C) with gentle agitation in PBS containing 0.3% Triton X-100 (T8787, Sigma-Aldrich), 0.5% bovine serum albumin (BSA; A2153, Sigma-Aldrich), and 1–10% normal donkey serum (AB7475, Abcam). Sections were then incubated overnight at 4°C in carrier solution (0.3% Triton X-100, 0.5% BSA, 1% normal donkey serum in PBS) containing primary antibodies: Rabbit polyclonal anti-GRIN1 (GluN1) (1:100–1:1000, #15141, BiCell Scientific), Goat polyclonal anti-ALDH1A1 (1:250–1:1000, #AF5869, R&D Systems), Chicken polyclonal anti-tyrosine hydroxylase (TH) (1:500–1:1000, #TYH, Aves Labs), Rabbit monoclonal anti-HA tag (1:500, #3724, Cell Signaling Technology).

After washing (3 × 10 min, PBS carrier solution), sections were incubated for 1 hour at room temperature with appropriate fluorescent donkey secondary antibodies conjugated to Alexa Fluor dyes (488, 546, 594, or 647; 1:500, Thermo Fisher Scientific), protected from light. Sections were then washed (2 × 10 min) and counterstained with DAPI (1:1000, 62248, Thermo Fisher Scientific) for 1 minute. After final PBS washes (2 × 10 min), sections were mounted on slides with ProLong® Gold antifade reagent (P36930, Invitrogen) and coverslipped.

Images were acquired on a Zeiss LSM 780 or 980 laser scanning confocal microscope using 20× and 63× (oil) objectives. Images were used for visualization of staining patterns and co-localization. For qualitative visualization, nonlinear lookup table (LUT) adjustments were applied; for quantitative analyses, no LUT modifications were made. Channel colors were standardized for presentation: TH (red), ALDH1A1 (blue), and GluN1 or HA (green).

### Quantification of CRISPR *Grin1* KD AAV transduction of ALDH1A1^+^ midbrain DANs

Consecutive 40 µm thick coronal midbrain sections spanning both the SNc and VTA were collected from viral *Grin1* KD mice (n = 2; left and right midbrain; one *Aldh1a1*^+/*P2A*-CreERT2^ mouse injected with AVV9-FLEX-SaCas9-U6-sg*Grin1*in the SNc, one *Aldh1a1*^+/*P2A*-CreERT2^ mouse injected with AVV9-FLEX-SaCas9-U6-sg*Grin1* in the VTA). Cells were stained with HA, ALDH1A1, TH, and DAPI. Images of the left and right midbrain from each section were taken with a laser scanning confocal microscope (LSM 980, Zeiss). HA^+^ and ALDH1A1^+^ cells were quantified using NeuroInfo (MBF Bioscience) using the “Detect Cells” pipeline. Regions of interest (ROIs) were manually drawn around the SNc and VTA using TH immunostaining and the Allen Brain Atlas as anatomical references. Channel-specific detection parameters were defined as follows: large diameter = 25.00 µm, small diameter = 8.00 µm, and detection strength = 1000. Following detection, colocalization analysis was conducted by using the “Colocalize Markers” pipeline. Quantification relied on the presence or absence of signal co-localization.

### mRNA sequencing

#### Experimental timeline pre-sample collection

Approximately 6-8-month-old female and male *Grin1* cKO and control mice were randomly assigned to undergo either 1) no food restriction (No Restriction group), 2) two consecutive nights of food restriction (Post-Restriction group), or 3) two consecutive nights of food restriction followed by three consecutive days of free access to food via an automatic feeding device (Refeeding group). Regardless of feeding condition, mice were moved from group housing to individual cages with access to their regular standard rodent chow and water. Mice in the No Restriction group were singly housed for approximately 48 hours before they were sacrificed, and their brains were collected. Mice in the Post-Restriction group were also sacrificed and their brains collected after approximately 48 hours, but food was removed from their cages overnight between the first and second day and the second and third day. Between the two overnight food restriction periods, standard rodent chow was put back into the cages. The mice in the Refeeding group underwent the same two nights of food restriction as the Post-Restriction group. However, after the second food restriction period, a FED3 device was added to each cage. The FED3 devices were set to “free feed” mode and loaded with 20 mg grain-based pellets (Dustless Precision Pellets®, #F0163, Bio-Serv). The mice then had constant free access to food via the feeding device for the next 72 hours. Each day at approximately 12:00 pm the devices were checked for jamming and pellet supplies were replenished. In the “free feed” mode, a pellet is automatically dispensed to a pellet receptor trough. If there is a pellet in the trough, another pellet will not be dispensed. Once the pellet is removed, the next pellet is automatically dispensed. After 72 hours of free access to food via the feeding device, the mice were sacrificed, and their brains were collected.

#### Sample collection

At the pre-determined experimental time points, mice were euthanized with CO_2_ followed by rapid decapitation. The brains were immediately dissected and split into two hemispheres. One hemisphere from each brain was placed in liquid nitrogen then stored at −80 °C. In total, 48 half brains were sent to AmpSeq for RNA extraction and mRNA sequencing (poly-A selection, 20 million reads). For each of the three different stages of food restriction (No Restriction, Post-Restriction, and Refeeding), 16 mice were collected: 4 male *Grin1* cKO mice, 4 male control mice, 4 female *Grin1* cKO mice, and 4 female control mice.

#### RNA extraction, raw data processing, and sequencing pipeline by AmpSeq

Raw sequencing reads were processed with fastp (v0.23.1) software to trim adapters and low-quality bases. The STAR (v2.7.11) aligner then mapped the trimmed reads to the mouse reference genome GRCm39. Next, the Salmon (v1.10.2) package was used to quantify transcript levels using a lightweight alignment-based method. Using the R package tximport (v1.30.0), a gene level summary of the transcript level quantification was created by mapping transcripts to genes and aggregating transcript abundance estimates.

#### Pre- and post-quality control by AmpSeq

Two different quality controls (QCs) were performed to identify any potential issues and biases in the trimmed and mapped reads. First, after initial fastp processing, the FastQC (v0.11.9) analysis tool was used to perform quality control to make sure that the trimmed reads were of high quality before proceeding. The MultiQC (v1.11) tool was then used to summarize the QC results across all samples in a single report. QC metrics for each trimmed sample, such as total number of sequences, sequence length range, GC percentage, total deduplicated percentage, and average sequence length were determined. Additional metrics determined included: sequence counts, sequence quality scores (mean per base and per sequence), per base sequence content (the proportion of each base position for which each of the four DNA bases had been called), per sequence GC percentage, per base N content, sequence length distribution, sequence duplication levels, amount of overrepresented sequences, and adapter content.

A second, post-alignment QC was performed using Qualimap (v2.3) to assess overall alignment quality. During this step, the quality of the aligned reads was assessed, information about the mapping distribution across different regions of the genome (e.g., exons, introns, and intergenic regions) was generated, and QC metrics such as coverage uniformity, strand specificity, and presence of PCR duplicates were determined.

#### Principal component analysis by AmpSeq

Principal component analysis (PCA) on Transcripts Per Million data was conducted to determine the main contributors of variation in the mRNA sequencing data.

#### Pearson correlation analysis

Pearson correlation analysis was used to assess the correlation strength between samples (including replicates) based on gene expression values.

#### Quantification by AmpSeq

Next, gene-level quantification was completed. Two different quantification measures were used. The number of reads that map to each gene (i.e., raw counts) was used to directly measure gene expression while Transcripts Per Million measurements accounted for different gene lengths and the number of mapped reads by normalizing the data. To determine Transcripts Per Million values, read counts were divided by gene length (kb) and then normalized to account for library size differences across samples.

#### mRNA sequencing differential gene expression analyses

Differential gene expression analyses were conducted using the DESeq2 R package [44] on raw counts at the gene level. PCAs done within the same sex and feeding group showed sample KC1, a male control mouse from the Refeeding group as an outlier. Going forward, it was excluded from groupwise comparisons. Pre-filtering excluded genes located on the X or Y chromosomes and any genes that were not expressed in more than half of the samples. The Negative Binomial Generalized Linear Model fitting and Wald statistics (nbinomWaldTest) were used to calculate log2 fold changes, Wald test p-values, and false discovery rate (FDR) adjusted p-values for each gene to compare *Grin1* cKO and control mice at each feeding stage, separately for males and females. The same model fitting and statistics were used to compare female and male mice at each feeding stage, separately for each genotype. Protein coding genes were considered significantly DEGs if their adjusted p-value was less than 0.05 and their log2 fold change was greater than 1.2. The DEGs were then analyzed using QIAGEN Ingenuity Pathway Analysis (QIAGEN Inc., https://digitalinsights.qiagen.com/IPA) to determine pathways that were significantly affected [45].

## Statistics

Data was analyzed using GraphPad Prism software (version 10.2.3). Specific statistical analyses and significant main effects of genotype are noted on the graphs of each figure. A p-value less than 0.05 was considered significant. (*p < 0.05, **p < 0.01, ***p <0.001, ****p < 0.0001).

## Inclusion & ethics statement

All experimental procedures were conducted in accordance with institutional guidelines and approved by the IACUC at NIA/NIH (protocol number: 464-LNG-2027). Every effort was made to minimize animal suffering and to reduce the number of animals used. Both male and female mice were included in all experiments, and sex was considered as a biological variable in experimental design and statistical analyses.

## Data availability

RNA-seq dataset have been deposited in the NCBI Gene Expression Omnibus (GEO) under accession number GSE309633 and are available at https://www.ncbi.nlm.nih.gov/geo/query/acc.cgi?acc=GSE309633.

## Code Availability

Not applicable.

## Funding Declaration

This work is supported in part by the Intramural Research Programs of National Institute on Aging, NIH (HC, ZIA AG000944, AG000928), and the Rodent Behavioral Core of Intramural Research Program of National Institute of Mental Health (MH002952). The contributions of the NIH author(s) were made as part of their official duties as NIH federal employees, are in compliance with agency policy requirements, and are considered Works of the United States Government. However, the findings and conclusions presented in this paper are those of the author(s) and do not necessarily reflect the views of the NIH or the U.S. Department of Health and Human Services.

## Acknowledgements

We thank Drs. Kate O’Connor-Giles, Zayd Khaliq, and Salinas Armando for their critical inputs, members of Cai lab, and the Rodent Behavioral Core of Intramural Research Program of National Institute of Mental Health for their suggestions and technical assistance.

## Author Contributions

H.C. conceived and wrote the manuscript and prepared the figures with inputs from all authors.

K.F.C. designed and performed mouse breeding, histology, behavioral experiments and data analyses, prepared figures, wrote methods and figure legends, and wrote the manuscript.

V.M.M.S. designed and conducted image analyses. J.D. performed gene expression analyses.

G.R. performed histology. L.C. performed stereotactic surgery and tissue dissection. L.S. built the control viral vector. L.W. designed FED3 experiments, contributed to data analysis, and edited the manuscript. All authors read and approved the final manuscript.

## Competing Interests

The authors declare no competing interests.

## Notes

### Competing Interest Statement

The authors have declared no competing interest.

## References

1. Jankovic J: Parkinson’s disease: clinical features and diagnosis. J Neurol Neurosurg Psychiatry 2008, 79(4):368–376.

2. de Chazeron I, Durif F, Chereau-Boudet I, Fantini ML, Marques A, Derost P, Debilly B, Brousse G, Boirie Y, Llorca PM: Compulsive eating behaviors in Parkinson’s disease. Eat Weight Disord 2019, 24(3):421–429.

3. Miwa H, Kondo T: Alteration of eating behaviors in patients with Parkinson’s disease: possibly overlooked? Neurocase 2008, 14(6):480–484.

4. de Chazeron I, Durif F, Lambert C, Chereau-Boudet I, Fantini ML, Marques A, Derost P, Debilly B, Brousse G, Boirie Y et al: A case-control study investigating food addiction in Parkinson patients. Sci Rep 2021, 11(1):10934.

5. Fearnley JM, Lees AJ: Ageing and Parkinson’s disease: substantia nigra regional selectivity. Brain 1991, 114 ( Pt 5):2283–2301.

6. Kordower JH, Olanow CW, Dodiya HB, Chu Y, Beach TG, Adler CH, Halliday GM, Bartus RT: Disease duration and the integrity of the nigrostriatal system in Parkinson’s disease. Brain 2013, 136(Pt 8):2419–2431.

7. Liu G, Yu J, Ding J, Xie C, Sun L, Rudenko I, Zheng W, Sastry N, Luo J, Rudow G et al: Aldehyde dehydrogenase 1 defines and protects a nigrostriatal dopaminergic neuron subpopulation. The Journal of clinical investigation 2014, 124(7):3032–3046.

8. Jacobs FM, Smits SM, Noorlander CW, von Oerthel L, van der Linden AJ, Burbach JP, Smidt MP: Retinoic acid counteracts developmental defects in the substantia nigra caused by Pitx3 deficiency. Development 2007, 134(14):2673–2684.

9. Kim JI, Ganesan S, Luo SX, Wu YW, Park E, Huang EJ, Chen L, Ding JB: Aldehyde dehydrogenase 1a1 mediates a GABA synthesis pathway in midbrain dopaminergic neurons. Science 2015, 350(6256):102–106.

10. Marchitti SA, Deitrich RA, Vasiliou V: Neurotoxicity and metabolism of the catecholamine-derived 3,4-dihydroxyphenylacetaldehyde and 3,4-dihydroxyphenylglycolaldehyde: the role of aldehyde dehydrogenase. Pharmacol Rev 2007, 59(2):125–150.

11. Galter D, Buervenich S, Carmine A, Anvret M, Olson L: ALDH1 mRNA: presence in human dopamine neurons and decreases in substantia nigra in Parkinson’s disease and in the ventral tegmental area in schizophrenia. Neurobiol Dis 2003, 14(3):637–647.

12. Martirosyan A, Ansari R, Pestana F, Hebestreit K, Gasparyan H, Aleksanyan R, Hnatova S, Poovathingal S, Marneffe C, Thal DR et al: Unravelling cell type-specific responses to Parkinson’s Disease at single cell resolution. Mol Neurodegener 2024, 19(1):7.

13. Pereira Luppi M, Azcorra M, Caronia-Brown G, Poulin JF, Gaertner Z, Gatica S, Moreno-Ramos OA, Nouri N, Dubois M, Ma YC et al: Sox6 expression distinguishes dorsally and ventrally biased dopamine neurons in the substantia nigra with distinctive properties and embryonic origins. Cell Rep 2021, 37(6):109975.

14. Wu J, Kung J, Dong J, Chang L, Xie C, Habib A, Hawes S, Yang N, Chen V, Liu Z et al: Distinct Connectivity and Functionality of Aldehyde Dehydrogenase 1a1-Positive Nigrostriatal Dopaminergic Neurons in Motor Learning. Cell Rep 2019, 28(5):1167–1181 e1167.

15. Crupi R, Impellizzeri D, Cuzzocrea S: Role of Metabotropic Glutamate Receptors in Neurological Disorders. Front Mol Neurosci 2019, 12:20.

16. Hunt DL, Castillo PE: Synaptic plasticity of NMDA receptors: mechanisms and functional implications. Curr Opin Neurobiol 2012, 22(3):496–508.

17. Carmichael K, Evans RC, Lopez E, Sun L, Kumar M, Ding J, Khaliq ZM, Cai H: Function and Regulation of ALDH1A1-Positive Nigrostriatal Dopaminergic Neurons in Motor Control and Parkinson’s Disease. Front Neural Circuits 2021, 15:644776.

18. Evans RC, Zhu M, Khaliq ZM: Dopamine Inhibition Differentially Controls Excitability of Substantia Nigra Dopamine Neuron Subpopulations through T-Type Calcium Channels. J Neurosci 2017, 37(13):3704–3720.

19. Zweifel LS, Parker JG, Lobb CJ, Rainwater A, Wall VZ, Fadok JP, Darvas M, Kim MJ, Mizumori SJ, Paladini CA et al: Disruption of NMDAR-dependent burst firing by dopamine neurons provides selective assessment of phasic dopamine-dependent behavior. Proc Natl Acad Sci U S A 2009, 106(18):7281–7288.

20. Wang LP, Li F, Wang D, Xie K, Wang D, Shen X, Tsien JZ: NMDA receptors in dopaminergic neurons are crucial for habit learning. Neuron 2011, 72(6):1055–1066.

21. Hutchison MA, Gu X, Adrover MF, Lee MR, Hnasko TS, Alvarez VA, Lu W: Genetic inhibition of neurotransmission reveals role of glutamatergic input to dopamine neurons in high-effort behavior. Mol Psychiatry 2018, 23(5):1213–1225.

22. Curtze C, Nutt JG, Carlson-Kuhta P, Mancini M, Horak FB: Levodopa Is a Double-Edged Sword for Balance and Gait in People With Parkinson’s Disease. Mov Disord 2015, 30(10):1361–1370.

23. Aradi SD, Hauser RA: Medical Management and Prevention of Motor Complications in Parkinson’s Disease. Neurotherapeutics 2020, 17(4):1339–1365.

24. Traynelis SF, Wollmuth LP, McBain CJ, Menniti FS, Vance KM, Ogden KK, Hansen KB, Yuan H, Myers SJ, Dingledine R: Glutamate receptor ion channels: structure, regulation, and function. Pharmacol Rev 2010, 62(3):405–496.

25. Monyer H, Sprengel R, Schoepfer R, Herb A, Higuchi M, Lomeli H, Burnashev N, Sakmann B, Seeburg PH: Heteromeric NMDA receptors: molecular and functional distinction of subtypes. Science 1992, 256(5060):1217–1221.

26. Cull-Candy SG, Leszkiewicz DN: Role of distinct NMDA receptor subtypes at central synapses. Sci STKE 2004, 2004(255):re16.

27. Schorge S, Colquhoun D: Studies of NMDA receptor function and stoichiometry with truncated and tandem subunits. J Neurosci 2003, 23(4):1151–1158.

28. Ulbrich MH, Isacoff EY: Subunit counting in membrane-bound proteins. Nat Methods 2007, 4(4):319–321.

29. Ulbrich MH, Isacoff EY: Rules of engagement for NMDA receptor subunits. Proc Natl Acad Sci U S A 2008, 105(37):14163–14168.

30. Paoletti P, Bellone C, Zhou Q: NMDA receptor subunit diversity: impact on receptor properties, synaptic plasticity and disease. Nat Rev Neurosci 2013, 14(6):383–400.

31. Tsien JZ, Chen DF, Gerber D, Tom C, Mercer EH, Anderson DJ, Mayford M, Kandel ER, Tonegawa S: Subregion- and cell type-restricted gene knockout in mouse brain. Cell 1996, 87(7):1317–1326.

32. Bonci A, Malenka RC: Properties and plasticity of excitatory synapses on dopaminergic and GABAergic cells in the ventral tegmental area. J Neurosci 1999, 19(10):3723–3730.

33. Overton PG, Richards CD, Berry MS, Clark D: Long-term potentiation at excitatory amino acid synapses on midbrain dopamine neurons. Neuroreport 1999, 10(2):221–226.

34. Ungless MA, Whistler JL, Malenka RC, Bonci A: Single cocaine exposure in vivo induces long-term potentiation in dopamine neurons. Nature 2001, 411(6837):583–587.

35. Prut L, Belzung C: The open field as a paradigm to measure the effects of drugs on anxiety-like behaviors: a review. Eur J Pharmacol 2003, 463(1-3):3–33.

36. Cousins MS, Salamone JD: Nucleus accumbens dopamine depletions in rats affect relative response allocation in a novel cost/benefit procedure. Pharmacol Biochem Behav 1994, 49(1):85–91.

37. Matikainen-Ankney BA, Earnest T, Ali M, Casey E, Wang JG, Sutton AK, Legaria AA, Barclay KM, Murdaugh LB, Norris MR et al: An open-source device for measuring food intake and operant behavior in rodent home-cages. Elife 2021, 10.

38. Hunker AC, Soden ME, Krayushkina D, Heymann G, Awatramani R, Zweifel LS: Conditional Single Vector CRISPR/SaCas9 Viruses for Efficient Mutagenesis in the Adult Mouse Nervous System. Cell Rep 2020, 30(12):4303–4316 e4306.

39. Cassidy RM, Tong Q: Hunger and Satiety Gauge Reward Sensitivity. Front Endocrinol (Lausanne) 2017, 8:104.

40. Habib A, Riccobono G, Tian L, Basu D, Sun L, Chang L, Martinez Smith VM, Wang L, Le W, Cai H: Subtype-Specific Roles of Nigrostriatal Dopaminergic Neurons in Motor and Associative Learning. bioRxiv 2025.

41. Hernandez L, Hoebel BG: Food reward and cocaine increase extracellular dopamine in the nucleus accumbens as measured by microdialysis. Life Sci 1988, 42(18):1705–1712.

42. Rada P, Avena NM, Hoebel BG: Daily bingeing on sugar repeatedly releases dopamine in the accumbens shell. Neuroscience 2005, 134(3):737–744.

43. Wilson C, Nomikos GG, Collu M, Fibiger HC: Dopaminergic correlates of motivated behavior: importance of drive. J Neurosci 1995, 15(7 Pt 2):5169–5178.

44. Love MI, Huber W, Anders S: Moderated estimation of fold change and dispersion for RNA-seq data with DESeq2. Genome Biol 2014, 15(12):550.

45. Kramer A, Green J, Pollard J, Jr., Tugendreich S: Causal analysis approaches in Ingenuity Pathway Analysis. Bioinformatics 2014, 30(4):523–530.

