## Supplementary material for "Sex-Dependent Effects of Glutamatergic Disruption on Dopaminergic Neuron Subtype Vulnerable in Parkinson’s Disease": Supplemtary files

### Supplementary Figures and Tables

**Supplementary Fig. 1**

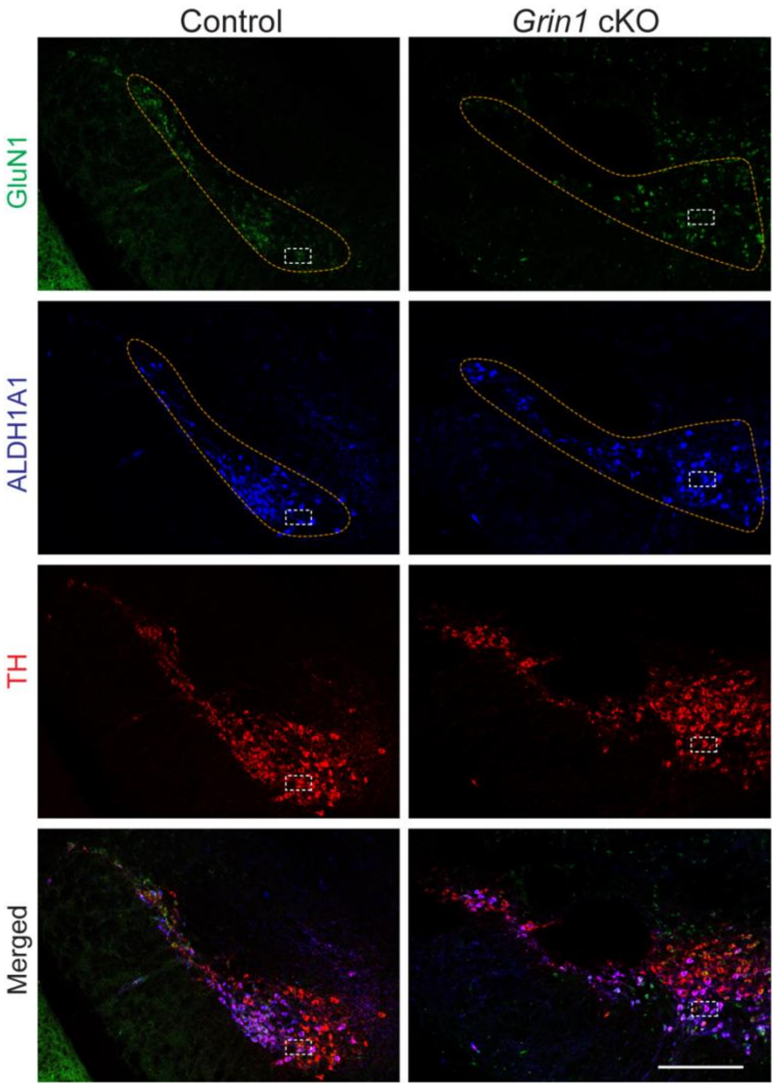


**Supplementary Fig. 1 Genetic deletion of *Grin1* in ALDH1A1[+] DANs.**

Representative image (20×, single z-plane) showing GluN1 (green), ALDH1A1 (blue), and TH (red) co-staining in *Grin1* cKO and control midbrain DANs outlined by orange dotted lines. White doted boxes indicate regions shown in Fig. 1b. Sale bar, 500 µm.

**Supplementary Fig. 2**

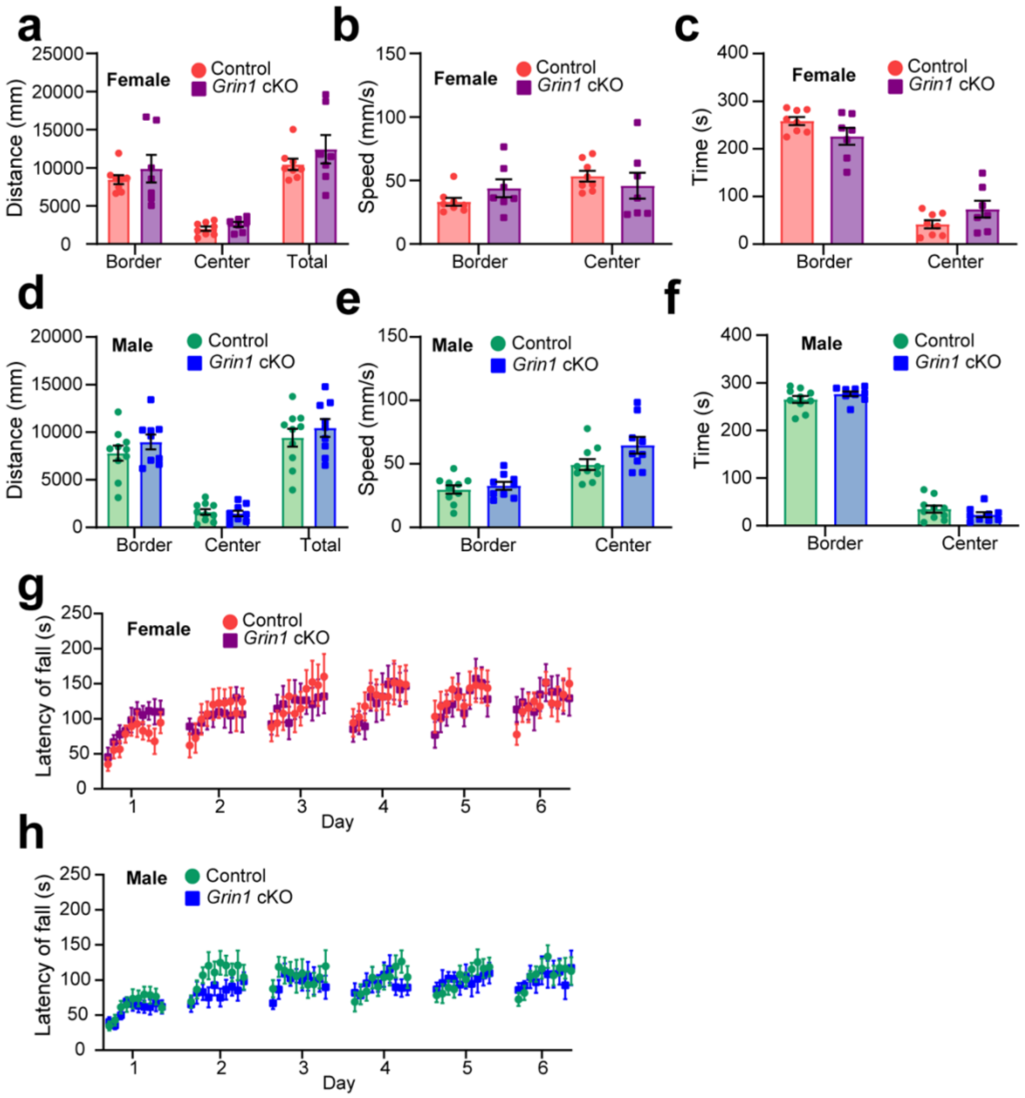


**Supplementary Fig. 2 No sex difference in spontaneous locomotion or motor learning.**

**a**, Distance traveled (mm) in the border, center, and total regions of the arena by female Grin1 cKO (n = 7) and control mice (n = 8). Unpaired t-tests: border, p = 0.4722; center, p = 0.2295; total, p = 0.3476.

**b**, Average speed (mm/s) in the border and center regions by female *Grin1* cKO (n = 7) and control mice (n = 8). Unpaired t-tests with Welch's correction: border, p = 0.2063; center, p = 0.5126.

**c**, Time spent (s) in the border and center of the open field arena by 3–4-month-old female *Grin1* cKO (n = 7) and control mice (n = 8) during a 5-min session. Unpaired t-tests with Welch's correction: border, p = 0.1382; center, p = 0.1385.

**d**, Distance traveled (mm) in border, center, and total regions by male *Grin1* cKO (n = 9) and control mice (n = 10). Unpaired t-tests with Welch's correction: border, p = 0.3085; center, p = 0.7143; total, p = 0.4490.

**e**, Average speed (mm/s) in border and center regions by male *Grin1* cKO (n = 9) and control mice (n = 10). Unpaired t-tests with Welch's correction: border, p = 0.5286; center, p = 0.0718.

**f**, Time spent (s) in border and center regions by 3–4-month-old male *Grin1* cKO (n = 9) and control mice (n = 10). Unpaired t-tests with Welch's correction: border, p = 0.2072; center, p = 0.2027.

**j**, Rotarod performance across 6 days (10 trials/day) in 3–4-month-old *Grin1* cKO (n = 7 females, 10 males) and control mice (n = 8 females, 12 males). Two-way ANOVA, genotype: F(1,35) = 0.7043, p = 0.7043.

**k**, Rotarod performance in female *Grin1* cKO (n = 7) and control mice (n = 8). Two-way ANOVA, genotype: F(1,13) = 0.0001432, p = 0.9906.

**l**, Rotarod performance in male *Grin1* cKO (n = 10) and control mice (n = 12). Two-way ANOVA, genotype:  F(1,20) = 0.4647, p = 0.5033.

All data are presented as mean ± SEM.

**Supplementary Fig. 3**

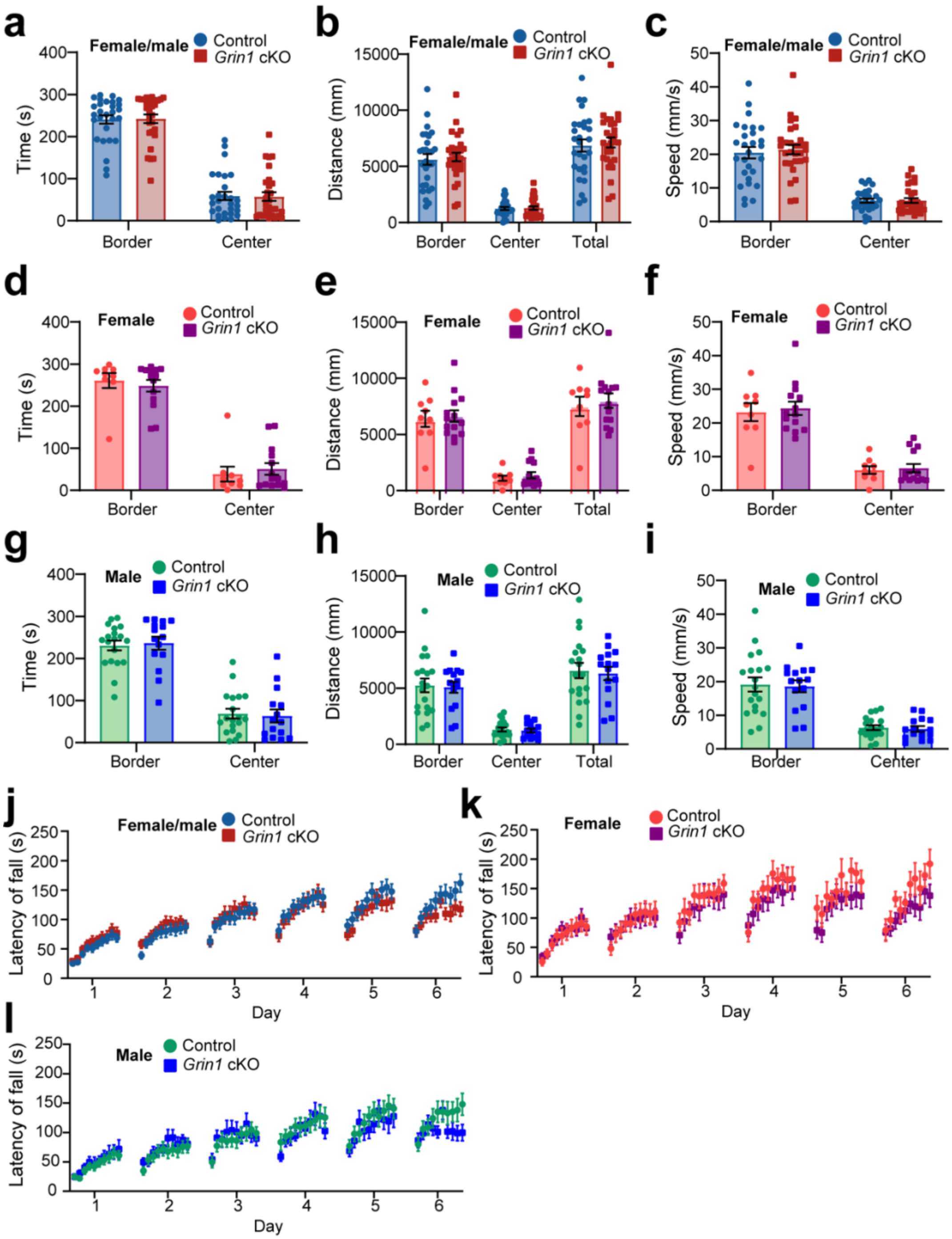


**Supplementary Fig. 3 Aged (15+ months) *Grin1* cKO mice display no deficits in spontaneous locomotion or motor learning.**

**a,** Time spent (s) in border and center regions of the open field arena by 15+ month-old (range 15–27 months) *Grin1* cKO (n = 14F, 15M) and control mice (n = 9F, 19M) during a 5-min session. Unpaired t-tests with Welch's correction: border, p = 0.9061; center, p = 0.9066.

**b**, Distance traveled (mm) in border, center, and total regions. Unpaired t-tests with Welch's correction: border, p = 0.7165; center, p = 0.8289; total, p = 0.7032.

**c**, Average speed (mm/s) in border and center regions. Unpaired t-tests with Welch's correction: border, p = 0.6788; center, p = 0.9750.

**d**, Time spent in border and center regions by aged female *Grin1* cKO (n = 14) and control (n = 9) mice. Unpaired t-tests with Welch's correction: border, p = 0.5873; center, p = 0.5873.

**e**, Distance traveled (mm) in border, center, and total regions by aged females. Unpaired t-tests with Welch's correction: border, p = 0.7827; center, p = 0.4689; total, p = 0.6400.

**f**, Average speed (mm/s) in border and center regions by aged females. Unpaired t-tests with Welch's correction: border, p = 0.7330; center, p = 0.7368.

**g**, Time spent in border and center regions by aged male *Grin1* cKO (n = 15) and control (n = 19) mice. Unpaired t-tests with Welch's correction: border, p = 0.7783; center, p = 0.7788.

**h**, Distance traveled (mm) in border, center, and total regions by aged males. Unpaired t-tests with Welch's correction: border, p = 0.8395; center, p = 0.7077; total, p = 0.7808.

**i**, Average velocity (mm/s) in border and center regions by aged males. Unpaired t-tests with Welch's correction: border, p = 0.8404; center, p = 0.7234.

**j**, Rotarod performance across 6 days (10 trials/day) in 15+ month-old *Grin1* cKO (n = 13F, 15M) and control mice (n = 9F, 20M). Two-way ANOVA, genotype: p = 0.8056.

**k**, Rotarod performance in aged females. Two-way ANOVA, genotype: p = 0.5185.

**l**, Rotarod performance in aged males. Two-way ANOVA, genotype: p = 0.8588.

All data are presented as mean ± SEM.

**Supplementary Fig. 4**

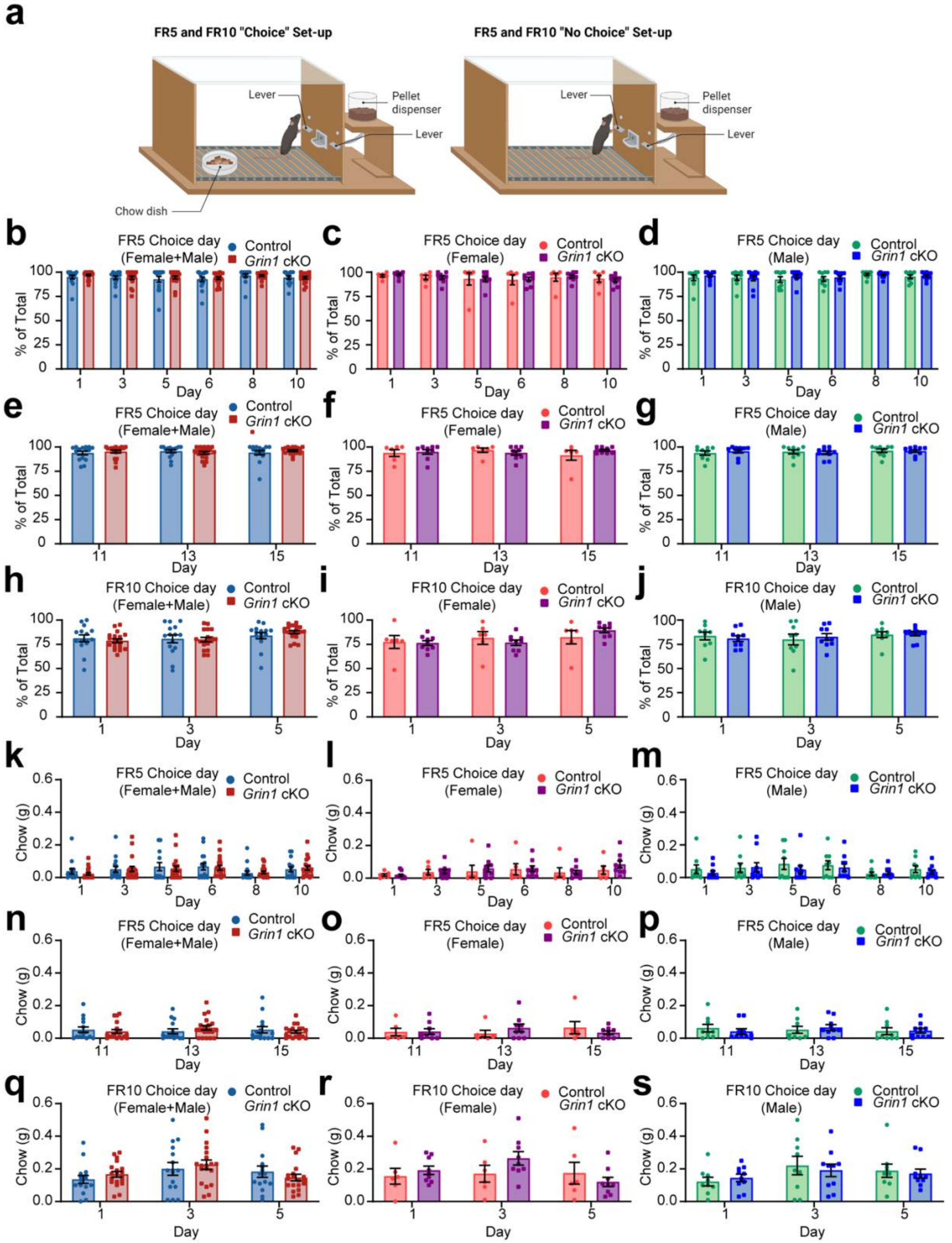

**Supplementary Fig. 4 Mice overwhelmingly choose to work for pellet rewards rather than consume freely available chow under high-effort choice conditions.**

**a**, Schematic of operant chamber setup for Choice and No Choice sessions, created by BioRender.

**b–d**, Percentage of total food consumed derived from chocolate pellet rewards on each FR5 Choice Day with a 50-pellet limit by Grin1 cKO (n = 9F, 10M) and control mice (n = 6F, 9M) for all mice (b), females only (c), and males only (d).

**e–g**, Percentage of total food consumed derived from chocolate pellet rewards on each FR5 Choice Day without a pellet limit for all mice (e), females only (f), and males only (g).

**h–j**, Percentage of total food consumed derived from chocolate pellet rewards on each FR10 Choice Day without a pellet limit for all mice (h), females only (i), and males only (j).

**k–m**, Chow consumed on each FR5 Choice Day with a 50-pellet limit for all mice (k), females only (l), and males only (m).

**n–p**, Chow consumed on each FR5 Choice Day without a pellet limit for all mice (n), females only (o), and males only (p).

**q–s**, Chow consumed on each FR10 Choice Day without a pellet limit for all mice (q), females only (r), and males only (s).

All data are presented as mean ± SEM.

**Supplementary Fig. 5**

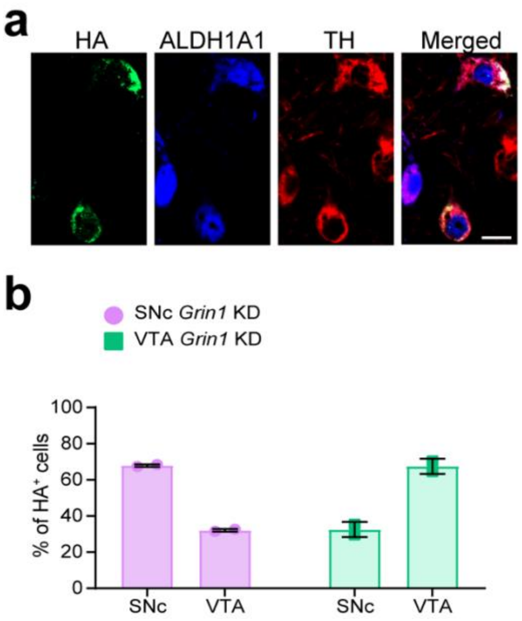


**Supplementary Fig. 5 Partial but regionally preferential knockdown of *Grin1* in SNc and VTA ALDH1A1$^+$ DANs.**

**a,** Representative confocal images (63×, single z-plane) showing HA (green), ALDH1A1 (blue), and TH (red) co-staining in midbrain coronal sections from *Aldh1a1*$^{+/P2A\text{-}CreERT2}$ mice injected with AAV9-FLEX-SaCas9-U6-sg*Grin1*. Scale bar, 20 µm.

**b**, Quantification of the percentage of HA$^+$ neurons among ALDH1A1$^+$ neurons in the SNc and VTA of *Grin1* KD mice. Data are presented as mean ± SEM.

**Supplementary Fig. 6**

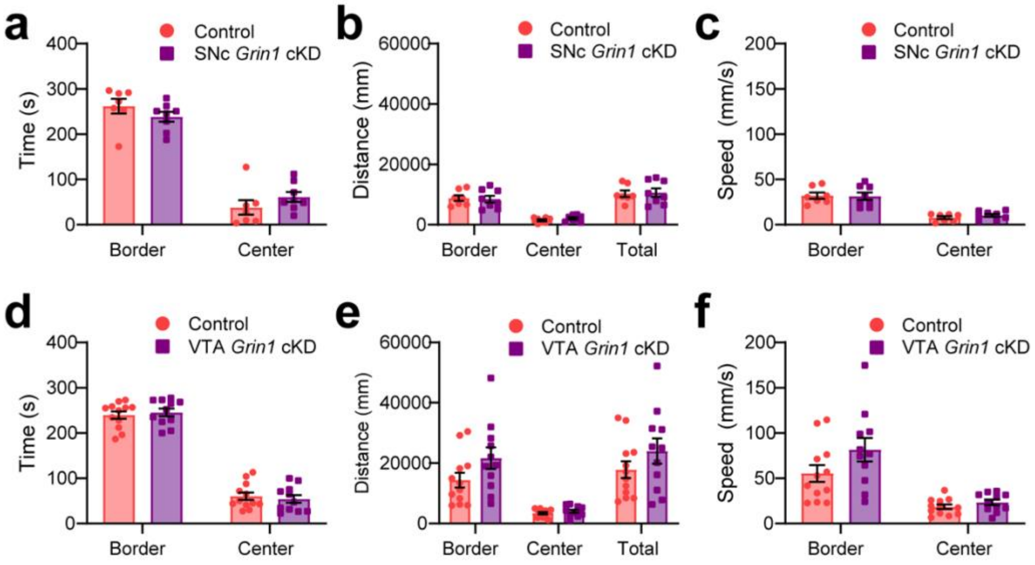


**Supplementary Fig. 6 Neither SNc nor VTA *Grin1* knockdown impairs spontaneous locomotion.**

**a–c,** Time spent (s) in border and center regions (a), distance traveled (mm) in border, center, and total regions (b), and average velocity (mm/s) (c) in female SNc *Grin1* KD mice (n = 8) and female controls (n = 7) during a 5-min open field session.

**d–f**, Time spent (s) in border and center regions (d), distance traveled (mm) in border, center, and total regions (e), and average velocity (mm/s) (f) in female VTA *Grin1* KD mice (n = 11) and female controls (n = 12) during a 5-min open field session.

All data are presented as mean ± SEM. Statistical comparisons were performed using unpaired t-tests. No significant changes were found.

**Supplementary Table 1 Female *Grin1* cKO vs. Control Post-Restriction (padj < 0.05, Fold Change (FC) > 1.2)**

| *GeneName* | *baseMean* | *log2FC* | *padj* |
|---|---|---|---|
| *H2-K1* | 274.048027 | 0.49638252 | 0.00640695 |
| *Col11a2* | 316.441569 | 0.45390244 | 0.03854266 |
| *Qdpr* | 2566.85077 | 0.45273172 | 4.68E-09 |
| *Mag* | 2146.60956 | 0.4473795 | 0.00640695 |
| *Col16a1* | 355.722652 | 0.43007071 | 0.03854266 |
| *Apoe* | 12023.6831 | 0.42703456 | 0.00012537 |
| *Cd63* | 905.118749 | 0.42293211 | 0.00221938 |
| *Igfbp2* | 305.407641 | 0.41292176 | 0.03579081 |
| *Baiap3* | 681.781968 | 0.40105077 | 0.01573579 |
| *Phgdh* | 622.164414 | 0.40072661 | 0.00221938 |
| *Gfap* | 2621.36072 | 0.3962146 | 0.04380223 |
| *Olig1* | 1103.20849 | 0.39371649 | 0.01512989 |
| *R3hcc1* | 280.236116 | 0.39143261 | 0.03060983 |
| *Pltp* | 503.965327 | 0.38862777 | 0.04249842 |
| *Itm2a* | 714.325178 | 0.38302008 | 0.01677498 |
| *Lpar1* | 471.072691 | 0.37922501 | 0.04333716 |
| *Guk1* | 1397.98688 | 0.37805874 | 0.04497013 |
| *Slc7a10* | 372.2885 | 0.36270342 | 0.02222805 |
| *Cmtm5* | 428.322447 | 0.35771038 | 0.04497013 |
| *Bloc1s1* | 526.122997 | 0.35492153 | 0.04333716 |
| *Cnp* | 3266.37051 | 0.34814951 | 0.00059299 |
| *Ndufb7* | 771.499959 | 0.34764064 | 0.03945405 |
| *H2-D1* | 616.357482 | 0.33041766 | 0.04380223 |
| *Phldb1* | 1365.46801 | 0.32373267 | 0.00640695 |
| *Mid1ip1* | 1092.51039 | 0.31629337 | 0.01524657 |
| *Pink1* | 5885.473 | 0.3075992 | 0.00059299 |
| *Bcan* | 2329.94115 | 0.30278552 | 0.01868968 |
| *Hnrnpa0* | 1269.29602 | 0.29554033 | 0.03060983 |
| *Ache* | 1062.43659 | 0.29458411 | 0.04333716 |
| *Cadm4* | 2330.2018 | 0.29012105 | 0.02222805 |
| *Sec11c* | 1018.76529 | 0.28584088 | 0.02300283 |
| *Bsg* | 3888.13154 | 0.27461338 | 0.00960625 |
| *Pum2* | 1867.35034 | -0.2671986 | 0.03854266 |
| *Cnot7* | 998.164703 | -0.2870596 | 0.04380223 |
| *Appl1* | 858.349384 | -0.2892618 | 0.01856791 |
| *Kcnj3* | 952.222791 | -0.289777 | 0.04984066 |

| | | | |
|---|---|---|---|
| *Ppm1e* | 1646.11248 | -0.2979408 | 0.04333716 |
| *Mtpn* | 3092.61337 | -0.2988075 | 0.04333716 |
| *Rock2* | 1378.07972 | -0.3131995 | 0.02237311 |
| *Far1* | 775.517807 | -0.314367 | 0.04096246 |
| *Pten* | 1670.37988 | -0.3236781 | 0.01652385 |
| *Ralgps2* | 410.00751 | -0.3281221 | 0.04380223 |
| *Spred1* | 1793.12301 | -0.3350164 | 0.00525725 |
| *Nrn1* | 1414.88207 | -0.3370176 | 0.01856791 |
| *Fut9* | 970.81522 | -0.353677 | 0.04333716 |
| *Bmi1* | 562.704336 | -0.3559021 | 0.04096246 |
| *Hook1* | 655.269096 | -0.3595119 | 0.01512989 |
| *Tut4* | 374.173951 | -0.3642117 | 0.03854266 |
| *Lrrtm2* | 573.647381 | -0.369953 | 0.01041116 |
| *Eeig2* | 560.46551 | -0.3722001 | 0.02802366 |
| *Kdm7a* | 561.548439 | -0.3798213 | 0.01652385 |
| *Nrip3* | 1658.79198 | -0.3808398 | 0.01652385 |
| *Zfp654* | 314.373061 | -0.3878257 | 0.04333716 |
| *Hycc2* | 915.786244 | -0.3930666 | 0.01652385 |
| *Mal2* | 1228.56137 | -0.394907 | 0.01380553 |
| *Ankrd45* | 546.709824 | -0.3954765 | 0.01780307 |
| *Fgd4* | 266.679158 | -0.396174 | 0.04497013 |
| *Tiam2* | 563.895236 | -0.4018757 | 0.01677498 |
| *Nlk* | 731.232006 | -0.4019342 | 0.00347561 |
| *Lrrc7* | 365.567793 | -0.4119464 | 0.01791048 |
| *Fam81a* | 1549.34139 | -0.4395466 | 0.00834767 |
| *Homer1* | 967.815243 | -0.458026 | 0.00083814 |
| *Ipcef1* | 608.751061 | -0.4773606 | 0.00655203 |
| *Clk1* | 1761.61205 | -0.5358466 | 0.01780307 |
| *Rnpc3* | 471.854253 | -0.5379202 | 0.00221938 |
| *Aldh1a1* | 1716.97002 | -0.753976 | 6.67E-06 |

**Supplementary Table 2 Female vs. Male *Grin1* cKO Post-Restriction (padj < 0.05, FC > 1.2)**

| *GeneName* | *baseMean* | *log2FC* | *padj* |
|---|---|---|---|
| *Opalin* | 199.874916 | 0.94074985 | 0.04999425 |
| *Mgp* | 161.580948 | 0.67253799 | 0.00039338 |
| *Cd59a* | 195.462801 | 0.61694186 | 0.00910119 |
| *Ppp1r14a* | 166.63343 | 0.60682339 | 0.00222927 |
| *Pdlim2* | 215.289431 | 0.55880949 | 0.00235 |
| *Pllp* | 504.51742 | 0.53079076 | 1.63E-05 |
| *Trf* | 3705.59642 | 0.51056199 | 2.62E-05 |
| *Fxyd1* | 189.142794 | 0.50412828 | 0.00692008 |
| *Qdpr* | 2390.4544 | 0.49835782 | 7.11E-10 |
| *Atp5me* | 1215.18914 | 0.49784423 | 0.00235 |
| *Snrnp25* | 196.175103 | 0.49224201 | 0.02784748 |
| *Uqcrh* | 2986.51328 | 0.48640747 | 0.00037395 |
| *Myo1d* | 163.166629 | 0.48124801 | 0.03320189 |
| *H1f2* | 247.668402 | 0.47866988 | 0.01202225 |
| *H2-K1* | 260.357738 | 0.47486901 | 0.00692008 |
| *Il33* | 644.619802 | 0.47231634 | 0.00222927 |
| *Mag* | 2012.45079 | 0.47015367 | 1.63E-05 |
| *Btbd17* | 161.53698 | 0.46365408 | 0.03581675 |
| *Gfap* | 2429.34713 | 0.45722079 | 0.02431769 |
| *Rps27* | 1885.49263 | 0.45612076 | 0.00079897 |
| *Tst* | 197.334377 | 0.4559967 | 0.0241288 |
| *Gstm7* | 200.506747 | 0.45370227 | 0.01533656 |
| *Cd63* | 846.298234 | 0.4511919 | 0.00011099 |
| *Mrpl2* | 275.828686 | 0.44476232 | 0.00609196 |
| *Rnaset2a* | 181.428401 | 0.44050004 | 0.02481636 |
| *Cd82* | 173.762047 | 0.42804489 | 0.04307917 |
| *Ndufb1* | 1010.6948 | 0.42603324 | 0.02061901 |
| *Anapc13* | 259.406356 | 0.42059922 | 0.03097916 |
| *Hebp1* | 249.814397 | 0.4175827 | 0.01667114 |
| *Phpt1* | 424.388478 | 0.41664707 | 0.02747186 |
| *Igfbp2* | 287.911073 | 0.41589941 | 0.01508212 |
| *C1qa* | 279.229178 | 0.41228435 | 0.008371 |
| *Ndufs5* | 926.533408 | 0.40866581 | 0.01358681 |
| *Penk* | 1564.37637 | 0.40123947 | 0.03911619 |
| *Cmtm5* | 398.878839 | 0.39606106 | 0.00707393 |
| *Syndig1l* | 465.000263 | 0.39255037 | 0.04205698 |
| *Ndufb7* | 717.729499 | 0.39158802 | 0.00141153 |

| | | | |
|---|---|---|---|
| *H2bc4* | 509.928065 | 0.39134133 | 0.02270128 |
| *Uqcr11* | 806.573092 | 0.38555179 | 0.00235 |
| *Bloc1s1* | 491.264356 | 0.38434903 | 0.00270135 |
| *Bola2* | 358.501684 | 0.38258403 | 0.02508383 |
| *Alg5* | 210.548904 | 0.38162031 | 0.04023602 |
| *Rpl36* | 1199.50634 | 0.38086685 | 0.03023662 |
| *Phlda3* | 263.437569 | 0.38007597 | 0.04585676 |
| *Bckdha* | 362.343364 | 0.37790632 | 0.03735992 |
| *Rpl38* | 2180.00411 | 0.37432007 | 0.00444585 |
| *Apoe* | 11527.3077 | 0.37267928 | 0.00141153 |
| *Atp5mf* | 1594.5351 | 0.37171686 | 0.03959799 |
| *Tmem256* | 601.267093 | 0.37150623 | 0.04310578 |
| *Cox6c* | 3831.05859 | 0.36866871 | 0.00222927 |
| *Rps29* | 2356.49297 | 0.36757092 | 0.00235 |
| *Mog* | 782.921298 | 0.36450358 | 0.00235 |
| *Guk1* | 1323.71794 | 0.36390309 | 0.03288297 |
| *Nme1* | 897.508809 | 0.36272577 | 0.00281137 |
| *Ctss* | 444.886186 | 0.36250566 | 0.01724156 |
| *Ddit4* | 1038.32883 | 0.35868748 | 0.02533006 |
| *Phgdh* | 595.266276 | 0.35631965 | 0.00235 |
| *Rps9* | 3090.92583 | 0.35586348 | 0.00316792 |
| *Cst3* | 8616.19169 | 0.35505611 | 0.00039338 |
| *Nudt18* | 316.984724 | 0.35238449 | 0.03248714 |
| *Naa38* | 318.677929 | 0.35185594 | 0.04347668 |
| *Mt3* | 2221.37983 | 0.35079473 | 0.00270135 |
| *Arsg* | 355.362921 | 0.34877974 | 0.03345851 |
| *mt-Nd4l* | 1742.82083 | 0.34760821 | 0.01190589 |
| *Cox7a2* | 2343.44043 | 0.34692908 | 0.02489946 |
| *Cnpy2* | 377.138635 | 0.34647336 | 0.02224116 |
| *S100a1* | 615.003761 | 0.34563186 | 0.01939294 |
| *Etfb* | 295.111922 | 0.345497 | 0.02784748 |
| *Ddrgk1* | 510.801159 | 0.34426685 | 0.00726593 |
| *Ndufa5* | 786.929057 | 0.34406408 | 0.00351926 |
| *Rps17* | 2332.01649 | 0.34380526 | 0.00107041 |
| *Ndufa2* | 653.057473 | 0.34278164 | 0.00476122 |
| *Slc35b2* | 376.533957 | 0.34269586 | 0.03693368 |
| *Rpl31* | 2286.49486 | 0.34167959 | 0.0057383 |
| *Rpl11* | 3273.06556 | 0.33983803 | 0.00848102 |
| *Plekhb1* | 7262.1324 | 0.33471872 | 0.04311491 |
| *Ndufa13* | 895.83493 | 0.33460388 | 0.02481636 |

| | | | |
|---|---|---|---|
| *Rpl37* | 1926.6503 | 0.33123862 | 0.04536576 |
| *Cldn10* | 344.244722 | 0.3309476 | 0.04999425 |
| *Naxd* | 419.628352 | 0.32935794 | 0.03959799 |
| *Olig1* | 1062.15109 | 0.32720673 | 0.03345851 |
| *Ramp1* | 832.150714 | 0.32644702 | 0.03133398 |
| *Rida* | 415.849514 | 0.32643231 | 0.04205698 |
| *S100b* | 2983.31911 | 0.32383379 | 0.04023602 |
| *Cldn11* | 1549.8037 | 0.32315673 | 0.00762478 |
| *Tmem63a* | 386.001098 | 0.32040423 | 0.04114355 |
| *Npc2* | 811.705375 | 0.31755809 | 0.02277991 |
| *Bsg* | 3618.39569 | 0.31721824 | 0.00039338 |
| *Ndufa3* | 1010.46075 | 0.31439631 | 0.01491784 |
| *Rpl36al* | 555.159784 | 0.31038991 | 0.03133398 |
| *Ndufb9* | 2909.26404 | 0.31033819 | 0.02067908 |
| *Sec11c* | 954.515685 | 0.31002765 | 0.04921266 |
| *Fau* | 2772.26924 | 0.30925653 | 0.04063552 |
| *Nnat* | 1657.96569 | 0.30776773 | 0.00075996 |
| *Slc7a10* | 356.959247 | 0.3064793 | 0.04205698 |
| *Gpr37* | 682.606424 | 0.30622763 | 0.01091217 |
| *Polr2e* | 767.347193 | 0.30469489 | 0.02127614 |
| *Tubb2b* | 799.66616 | 0.3008951 | 0.00955448 |
| *Rps8* | 3970.91495 | 0.29818337 | 0.00242693 |
| *Rps28* | 1109.56787 | 0.29683404 | 0.04336012 |
| *Cox5a* | 1086.22385 | 0.29497217 | 0.0124189 |
| *Cnp* | 3134.58024 | 0.29435216 | 0.00051371 |
| *Ptn* | 2236.2406 | 0.29408069 | 0.00716801 |
| *Mrps33* | 766.54243 | 0.29170123 | 0.03735992 |
| *Hcn2* | 2172.60828 | 0.28968612 | 0.03459921 |
| *Uqcrq* | 1591.38384 | 0.28817619 | 0.03133398 |
| *Rasgrp2* | 568.888709 | 0.28802514 | 0.03459885 |
| *Clta* | 1640.09319 | 0.28617846 | 0.01192743 |
| *Mesd* | 542.762661 | 0.28588984 | 0.03133398 |
| *Tmem50a* | 986.570912 | 0.28439376 | 0.01190589 |
| *Gpr37l1* | 2268.0289 | 0.28403024 | 0.00647163 |
| *Prmt1* | 692.261807 | 0.2818083 | 0.02747186 |
| *Mrps7* | 502.806085 | 0.27886086 | 0.04023602 |
| *Capns1* | 886.428581 | 0.27457747 | 0.0241288 |
| *Car2* | 1792.1873 | 0.27438941 | 0.00278746 |
| *Cd81* | 2595.37801 | 0.27016593 | 0.0124189 |
| *Aldh2* | 944.141597 | 0.26981238 | 0.03735992 |

| | | | |
|---|---|---|---|
| *Rpl37a* | 2226.16943 | 0.2676973 | 0.04473761 |
| *Arl6ip4* | 487.480269 | 0.26642822 | 0.04999425 |
| *Psmb4* | 1377.15057 | 0.26642703 | 0.04023602 |
| *Fabp5* | 996.42283 | 0.26638222 | 0.04660039 |
| *Snf8* | 578.652235 | 0.26323149 | 0.03832747 |
| *Spcs2* | 787.227968 | 0.26141906 | 0.03345851 |
| *Rpl19* | 4501.14433 | 0.26128353 | 0.02429898 |
| *Nktr* | 1447.89973 | -0.2605061 | 0.00703059 |
| *Spred1* | 1643.92332 | -0.2624098 | 0.03832747 |
| *Plxna2* | 987.325326 | -0.2638824 | 0.03074449 |
| *Dusp11* | 826.223855 | -0.2639099 | 0.03965821 |
| *Zfp207* | 1233.62465 | -0.2653552 | 0.0241288 |
| *Nvl* | 716.962874 | -0.2657045 | 0.03680651 |
| *Chl1* | 925.408465 | -0.2666516 | 0.04454387 |
| *Ylpm1* | 877.332431 | -0.2715057 | 0.04742388 |
| *Fryl* | 888.522594 | -0.2715708 | 0.02429898 |
| *Usp34* | 1293.22979 | -0.2716007 | 0.03911619 |
| *Ppm1e* | 1538.46219 | -0.2735011 | 0.02330335 |
| *Nrsn1* | 5406.05617 | -0.2752976 | 0.03028327 |
| *Lyst* | 749.05937 | -0.2761712 | 0.02508383 |
| *Gria2* | 3798.83189 | -0.2781645 | 0.00910119 |
| *Nfat5* | 853.577623 | -0.2822747 | 0.03959799 |
| *Nrip3* | 1506.97101 | -0.284056 | 0.03735992 |
| *Rock2* | 1286.69354 | -0.2871329 | 0.04473761 |
| *Ankhd1* | 628.884585 | -0.287384 | 0.04584398 |
| *Vps37a* | 664.661711 | -0.2901274 | 0.04702466 |
| *Kmt2a* | 1808.46748 | -0.2910644 | 0.02434852 |
| *Dennd5b* | 666.886824 | -0.2936928 | 0.04311491 |
| *Tnrc6b* | 871.033382 | -0.2956406 | 0.02840349 |
| *Etv1* | 1313.89176 | -0.2959571 | 0.04585676 |
| *Med12l* | 451.797515 | -0.2976669 | 0.03097916 |
| *Mkln1* | 830.841138 | -0.2995261 | 0.02481636 |
| *Cpeb4* | 1004.12027 | -0.2996537 | 0.04023602 |
| *Mga* | 670.533905 | -0.2997963 | 0.03684423 |
| *Rora* | 937.072956 | -0.2998707 | 0.01909128 |
| *Mapk8* | 847.300897 | -0.3042103 | 0.03959799 |
| *Zfc3h1* | 494.069482 | -0.3042639 | 0.04702466 |
| *Tlcd4* | 495.712597 | -0.3046329 | 0.03959799 |
| *Nup153* | 484.770096 | -0.3059496 | 0.02543527 |
| *Hook3* | 687.876442 | -0.3082905 | 0.04999425 |

| | | | |
|---|---|---|---|
| *Shprh* | 472.057525 | -0.3089192 | 0.03798956 |
| *Arfgef3* | 1155.92715 | -0.3101574 | 0.02316161 |
| *Ankrd45* | 498.129284 | -0.3104534 | 0.04311491 |
| *Tnr* | 1136.06405 | -0.3125643 | 0.00703059 |
| *Pgm2l1* | 1928.85845 | -0.3131839 | 0.02508383 |
| *Ipcef1* | 537.175608 | -0.3141042 | 0.03097916 |
| *Cacnb4* | 1841.38136 | -0.3157404 | 0.01419834 |
| *Dmxl1* | 514.551682 | -0.3166876 | 0.02508383 |
| *Hipk3* | 999.120306 | -0.3174596 | 0.00968437 |
| *Cep170* | 679.129646 | -0.317933 | 0.03074449 |
| *Rabgap1l* | 901.391667 | -0.3192435 | 0.01514674 |
| *Tank* | 328.396257 | -0.3193817 | 0.04765109 |
| *Cpsf6* | 847.281986 | -0.3224825 | 0.02067908 |
| *Braf* | 1041.80927 | -0.3233377 | 0.02508383 |
| *Slf2* | 798.513102 | -0.324079 | 0.00430383 |
| *Nlk* | 668.452987 | -0.3242851 | 0.02326657 |
| *Far1* | 734.712724 | -0.324944 | 0.02316161 |
| *Wsb1* | 1381.79811 | -0.3252097 | 0.04805834 |
| *Ubr3* | 1731.155 | -0.3271569 | 0.00463407 |
| *Tnpo1* | 644.998921 | -0.3272968 | 0.02747186 |
| *Ankrd12* | 639.106482 | -0.3290678 | 0.01202225 |
| *Rictor* | 642.979138 | -0.3322807 | 0.01688371 |
| *Kmt2c* | 1034.08669 | -0.3351662 | 0.00968437 |
| *Vps13a* | 385.939452 | -0.3365287 | 0.04063552 |
| *Pik3c2a* | 334.979461 | -0.3368655 | 0.04311491 |
| *Fut9* | 910.497928 | -0.3384678 | 0.04111873 |
| *Sfpq* | 1252.16807 | -0.3453776 | 0.0114219 |
| *Hook1* | 615.099329 | -0.3454017 | 0.01194294 |
| *Prkaa2* | 335.683314 | -0.3471655 | 0.03063038 |
| *Lrrtm2* | 538.335185 | -0.3529534 | 0.03345851 |
| *Nipbl* | 477.845127 | -0.3537171 | 0.01709692 |
| *Nrip1* | 326.917246 | -0.3557632 | 0.02234918 |
| *Fam81a* | 1412.77688 | -0.3568133 | 0.02747186 |
| *Zfp292* | 309.498182 | -0.3571794 | 0.03133398 |
| *Atp11b* | 1229.19529 | -0.357187 | 0.02747186 |
| *Dcp1a* | 236.645561 | -0.3625635 | 0.04454387 |
| *Rfx3* | 330.423167 | -0.3639236 | 0.02747186 |
| *Ppargc1a* | 458.645812 | -0.3656208 | 0.01724156 |
| *Fam135b* | 514.962848 | -0.3779701 | 0.02434852 |
| *Acvr2a* | 408.123628 | -0.3822578 | 0.03067977 |

| | | | |
|---|---|---|---|
| *Faxc* | 371.918185 | -0.3849505 | 0.01383436 |
| *Homer2* | 468.505728 | -0.3868798 | 0.01839618 |
| *Hipk2* | 638.732817 | -0.3882349 | 0.03735992 |
| *Tia1* | 1296.18244 | -0.3971213 | 0.00235 |
| *Exph5* | 252.689421 | -0.4129747 | 0.02481636 |
| *Gng4* | 394.572471 | -0.4162777 | 0.02449833 |
| *Smim43* | 286.805525 | -0.4208916 | 0.04753153 |
| *Phip* | 468.698432 | -0.4221888 | 0.00848102 |
| *Clk1* | 1590.18256 | -0.4271787 | 0.00968437 |
| *Klf12* | 172.484175 | -0.4298338 | 0.04205698 |
| *Lin7a* | 277.659033 | -0.4313439 | 0.02854909 |
| *Tut4* | 362.868084 | -0.4327049 | 0.00235 |
| *Fgd4* | 257.539898 | -0.4537412 | 0.00903222 |
| *Ndst3* | 351.639683 | -0.4674083 | 0.00256019 |
| *Lcorl* | 186.762233 | -0.4714637 | 0.04023602 |
| *Rnpc3* | 437.030373 | -0.4931622 | 0.00235 |
| *Dlx1* | 184.713189 | -0.49838 | 0.04999425 |
| *Hectd2* | 158.600225 | -0.5469694 | 0.00707393 |
| *Stxbp5l* | 317.683406 | -0.5915485 | 0.00059286 |

**Supplementary Table 3 Female vs. Male *Grin1* cKO Refeeding (padj < 0.05, FC > 1.2)**

| *GeneName* | *baseMean* | *log2FC* | *padj* |
|---|---|---|---|
| *Crtam* | 159.218579 | 0.89397572 | 0.00022375 |
| *Ppp1r17* | 253.988641 | 0.72578874 | 0.00200338 |
| *Car8* | 2498.4762 | 0.72240612 | 0.02547201 |
| *Pvalb* | 1846.61872 | 0.65055657 | 2.15E-07 |
| *Cd59a* | 171.043912 | 0.63824539 | 0.00234615 |
| *Rpa3* | 154.101342 | 0.60316832 | 0.01127369 |
| *Sar1b* | 813.837586 | 0.59885807 | 0.00083282 |
| *Gm56450* | 422.422412 | 0.54903475 | 0.00268575 |
| *Nrep* | 1843.68119 | 0.54179683 | 0.00087596 |
| *Selenof* | 2470.11471 | 0.53837754 | 0.00022375 |
| *Sumo1* | 1798.66746 | 0.51711939 | 0.00088785 |
| *Ciao2a* | 443.934127 | 0.51378241 | 0.00705974 |
| *Smim26* | 296.23253 | 0.5129004 | 0.01483541 |
| *Mkks* | 204.729255 | 0.50741723 | 0.00359634 |
| *Ggh* | 225.183061 | 0.50693699 | 0.01367739 |
| *Cox16* | 225.809947 | 0.50303026 | 0.02331741 |
| *Tm2d1* | 543.3849 | 0.5026828 | 0.00922603 |
| *Cep15* | 329.349667 | 0.49816659 | 0.00248676 |
| *Smim8* | 174.196141 | 0.49782562 | 0.03479579 |
| *Ube2v2* | 1412.36935 | 0.49709547 | 0.04375039 |
| *Pigx* | 181.337623 | 0.49620722 | 0.03563207 |
| *Acyp1* | 338.912758 | 0.49077026 | 0.0092212 |
| *Haus1* | 123.351236 | 0.48798978 | 0.01844832 |
| *Ccdc90b* | 260.427153 | 0.48791754 | 0.00524867 |
| *Gng11* | 221.121133 | 0.48706103 | 0.01181282 |
| *Ndufs4* | 1317.49169 | 0.48599827 | 0.00087596 |
| *Medag* | 259.081283 | 0.48538178 | 0.00093949 |
| *Gng13* | 459.923368 | 0.48324629 | 0.00456836 |
| *1110059E24Rik* | 379.561152 | 0.47178511 | 0.01844832 |
| *A430005L14Rik* | 229.731683 | 0.47166086 | 0.0226924 |
| *Zfp560* | 122.160199 | 0.46959317 | 0.03584175 |
| *Stk17b* | 125.017982 | 0.46771851 | 0.02990791 |
| *Rpp30* | 148.576706 | 0.46750802 | 0.01483541 |
| *Cript* | 764.976106 | 0.46585894 | 0.0033047 |
| *Hypk* | 171.056194 | 0.46238356 | 0.03414936 |
| *Lyz2* | 208.628607 | 0.4623632 | 0.03562166 |
| *1810037I17Rik* | 819.777108 | 0.45948579 | 0.00308567 |

| | | | |
|---|---|---|---|
| *Smim15* | 671.155631 | 0.45876694 | 0.00248676 |
| *Mien1* | 610.301249 | 0.45777964 | 0.00142688 |
| *C330018D20Rik* | 189.314933 | 0.45718468 | 0.02415697 |
| *Cox7c* | 2204.17876 | 0.45675723 | 0.00200338 |
| *Slirp* | 405.851958 | 0.45632079 | 0.0212338 |
| *Psma5* | 1152.31907 | 0.45422354 | 0.00532254 |
| *Tmem126a* | 255.366978 | 0.45405048 | 0.04230427 |
| *C1d* | 736.283195 | 0.45086345 | 0.01558959 |
| *Arl6* | 285.523504 | 0.45072207 | 0.00248676 |
| *Tma7* | 1327.7082 | 0.44907872 | 0.00456836 |
| *Eef1e1* | 287.557117 | 0.44831658 | 0.00560632 |
| *Zbtb26* | 129.455755 | 0.44768482 | 0.02239612 |
| *Itgb1bp1* | 465.920718 | 0.44602269 | 0.01090116 |
| *Gm2007* | 193.921486 | 0.44583367 | 0.02529826 |
| *Ndufaf4* | 400.84964 | 0.44485018 | 0.00292376 |
| *Etfrf1* | 366.302934 | 0.44456897 | 0.0054127 |
| *Mrpl14* | 258.52657 | 0.44374077 | 0.01782609 |
| *Nup35* | 179.643367 | 0.44059955 | 0.02565907 |
| *Mrpl20* | 731.915987 | 0.43900275 | 0.00429778 |
| *Ndufb6* | 1147.40063 | 0.43825918 | 0.00456836 |
| *Ccdc126* | 137.873245 | 0.43815097 | 0.01795213 |
| *Nts* | 142.939001 | 0.43592648 | 0.03847688 |
| *Zfp874a* | 182.540693 | 0.4352959 | 0.01441162 |
| *Med21* | 362.104708 | 0.4349063 | 0.01127369 |
| *Eif3e* | 935.98704 | 0.4345202 | 0.00650312 |
| *Sptssa* | 492.682882 | 0.43396571 | 0.02645887 |
| *Bcap29* | 568.697009 | 0.43370519 | 0.00256791 |
| *Cebpzos* | 286.796739 | 0.43353978 | 0.01777441 |
| *Gabarapl2* | 3885.27501 | 0.4326535 | 0.00090963 |
| *Glmn* | 169.870204 | 0.43222724 | 0.01092879 |
| *Tnfaip6* | 176.417534 | 0.43213772 | 0.01827321 |
| *Gm10033* | 214.413761 | 0.43192727 | 0.01368721 |
| *Timp4* | 822.324868 | 0.43113181 | 0.00222097 |
| *Lamtor5* | 686.511083 | 0.43064319 | 0.00892933 |
| *Lsm3* | 204.582162 | 0.43016525 | 0.01514005 |
| *Ift20* | 806.802467 | 0.42696983 | 0.00452523 |
| *Pde5a* | 176.457537 | 0.42575551 | 0.03666908 |
| *Acyp2* | 373.127797 | 0.42464294 | 0.02415697 |
| *Nop10* | 584.236908 | 0.42463627 | 0.01476885 |
| *Cops9* | 1380.97414 | 0.42360315 | 0.0082968 |

| | | | |
|---|---|---|---|
| *Amn1* | 176.988828 | 0.42273631 | 0.02897421 |
| *Ostf1* | 208.690729 | 0.41871075 | 0.01090116 |
| *Med6* | 229.704434 | 0.41823943 | 0.01777441 |
| *Rpl24* | 2066.71519 | 0.41757455 | 0.00136166 |
| *Cetn3* | 733.928284 | 0.41755636 | 0.00705974 |
| *Cstb* | 289.097502 | 0.41701453 | 0.02298864 |
| *Ndufa7* | 1224.30585 | 0.41668021 | 0.00952846 |
| *Clybl* | 242.093748 | 0.41581342 | 0.01302628 |
| *Sdhaf4* | 297.866665 | 0.41537224 | 0.03703514 |
| *Prdx1* | 1920.88334 | 0.41428575 | 0.00827754 |
| *Cnih1* | 938.586799 | 0.41351181 | 0.00313367 |
| *Ap3s1* | 576.247138 | 0.41345369 | 0.00369641 |
| *Neurod1* | 600.583283 | 0.41306103 | 0.03032892 |
| *Elof1* | 647.76233 | 0.41047047 | 0.01661174 |
| *Tmem60* | 244.226434 | 0.40872778 | 0.01181282 |
| *Commd6* | 470.862519 | 0.40819977 | 0.02483492 |
| *Mtln* | 343.088648 | 0.40792099 | 0.01068415 |
| *Mrps21* | 379.412961 | 0.406916 | 0.0092212 |
| *Rnf138* | 168.909236 | 0.4067512 | 0.02395583 |
| *Mrps14* | 361.788696 | 0.40557257 | 0.00705974 |
| *Tomm5* | 515.828915 | 0.40514023 | 0.00200338 |
| *Ostc* | 430.031465 | 0.4048656 | 0.0251625 |
| *Vamp8* | 124.351381 | 0.40274332 | 0.04542842 |
| *Mrpl33* | 406.279793 | 0.40130045 | 0.03502705 |
| *Vps29* | 1934.91502 | 0.40122281 | 0.00222097 |
| *Dph3* | 555.546808 | 0.40120073 | 0.00862459 |
| *Alkbh7* | 201.127463 | 0.40118599 | 0.04629228 |
| *Mrpl34* | 154.59011 | 0.4007841 | 0.04296245 |
| *Mrpl13* | 473.240494 | 0.40036263 | 0.01022591 |
| *Psmc6* | 1158.66011 | 0.40003435 | 0.00827754 |
| *Commd1* | 282.932957 | 0.39905922 | 0.00862459 |
| *Fcf1* | 233.966688 | 0.39625636 | 0.0146222 |
| *Eif4a2* | 11679.7303 | 0.39536538 | 0.00202836 |
| *H3f3a* | 2507.32916 | 0.39400986 | 0.01441162 |
| *Sec61g* | 673.219848 | 0.39372494 | 0.03562166 |
| *Spryd7* | 668.792657 | 0.39131381 | 0.0046996 |
| *Dusp19* | 238.75865 | 0.39093097 | 0.03465087 |
| *Hopx* | 820.126682 | 0.39005186 | 0.02072567 |
| *Rpl35a* | 3012.92519 | 0.38922381 | 0.01483541 |
| *Gng10* | 581.453629 | 0.38871657 | 0.01368721 |

| | | | |
|---|---|---|---|
| *Ndufc2* | 1313.36515 | 0.38530469 | 0.00569148 |
| *Eif2s2* | 1085.85455 | 0.3836793 | 0.00452523 |
| *Mreg* | 166.61177 | 0.38311541 | 0.04542842 |
| *Mob4* | 723.321762 | 0.38265528 | 0.0263376 |
| *Bnip3* | 1712.04543 | 0.38198338 | 0.00359634 |
| *Dbi* | 2005.46447 | 0.38155932 | 0.0161822 |
| *Bbip1* | 592.210771 | 0.38092237 | 0.04625252 |
| *Sav1* | 170.436827 | 0.38087356 | 0.03230583 |
| *Cryzl1* | 889.07824 | 0.38002843 | 0.00387091 |
| *Bnip3l* | 2220.90116 | 0.37868501 | 0.00093949 |
| *Chpt1* | 914.081217 | 0.37818535 | 0.00234615 |
| *Sdhaf3* | 218.447251 | 0.37668692 | 0.0256611 |
| *Snap23* | 217.337079 | 0.37616719 | 0.0294758 |
| *Septin7* | 4576.53842 | 0.37511218 | 0.00034827 |
| *Zfp874b* | 202.017854 | 0.37455319 | 0.03666908 |
| *Gm14325* | 242.011731 | 0.37432275 | 0.03470854 |
| *Naa20* | 697.621987 | 0.37419101 | 0.00862459 |
| *Cops3* | 665.420873 | 0.37384234 | 0.00369641 |
| *Hspe1* | 936.64685 | 0.37304637 | 0.02790403 |
| *Dctn3* | 980.63447 | 0.3715825 | 0.00433875 |
| *Crls1* | 366.812584 | 0.37093937 | 0.02790403 |
| *Ost4* | 406.461724 | 0.3704957 | 0.01110346 |
| *Polb* | 453.205696 | 0.36999242 | 0.00237524 |
| *Mrpl41* | 765.946499 | 0.36852729 | 0.03841817 |
| *Naca* | 2830.21904 | 0.36835479 | 0.01155129 |
| *Chrac1* | 199.999496 | 0.3683459 | 0.03045413 |
| *Tmed5* | 280.616309 | 0.36769903 | 0.00859357 |
| *Rtraf* | 831.971282 | 0.36767826 | 0.03281345 |
| *Uqcc2* | 1072.67192 | 0.36511761 | 0.01777441 |
| *Emc6* | 438.062871 | 0.36500508 | 0.00859357 |
| *Rap1a* | 662.166471 | 0.36457736 | 0.00271299 |
| *Coa6* | 213.233474 | 0.36323583 | 0.04324808 |
| *Rps20* | 1955.15052 | 0.36124318 | 0.0212338 |
| *Ndufc1* | 536.208685 | 0.36114638 | 0.04243815 |
| *Tmem242* | 774.198097 | 0.35929727 | 0.0054127 |
| *S100a1* | 589.736293 | 0.35823551 | 0.02089038 |
| *Ifrd1* | 433.595935 | 0.35714727 | 0.0161822 |
| *Scoc* | 1662.5468 | 0.35674584 | 0.00170948 |
| *Sec22a* | 188.847681 | 0.35631277 | 0.03179971 |
| *Erh* | 751.250896 | 0.3556153 | 0.01782609 |

| | | | |
|---|---|---|---|
| *Gnpda2* | 222.008057 | 0.35419774 | 0.03840906 |
| *Cln5* | 238.588931 | 0.35342828 | 0.02747102 |
| *Arl1* | 1961.82501 | 0.35200315 | 0.00165976 |
| *Uqcrq* | 1378.36687 | 0.35197737 | 0.01239244 |
| *Mrps33* | 727.451692 | 0.35129816 | 0.01558959 |
| *Lamtor3* | 532.662263 | 0.35093923 | 0.0092212 |
| *Btg1* | 557.221075 | 0.35021081 | 0.02498746 |
| *Fkbp3* | 1777.47498 | 0.35019076 | 0.00136166 |
| *Rps10* | 2549.91638 | 0.35008069 | 0.02581572 |
| *Commd8* | 592.750761 | 0.34975845 | 0.01023318 |
| *Aspa* | 276.407434 | 0.34974973 | 0.02236469 |
| *Ccng1* | 838.028612 | 0.34951854 | 0.00268553 |
| *Nck1* | 252.84717 | 0.34946137 | 0.02635808 |
| *Nsmce2* | 222.815654 | 0.34933072 | 0.04314042 |
| *Umad1* | 511.309007 | 0.34797843 | 0.01155129 |
| *Skp1* | 5446.5927 | 0.34733267 | 0.00452523 |
| *Mrpl21* | 258.514007 | 0.34634112 | 0.02793778 |
| *Csgalnact2* | 226.960003 | 0.34632539 | 0.01912429 |
| *Grpel1* | 579.656278 | 0.3460206 | 0.00696551 |
| *Selenok* | 1164.92445 | 0.34585698 | 0.0092212 |
| *Sf3b6* | 616.198644 | 0.34549927 | 0.04593503 |
| *Lcorl* | 189.884626 | 0.34531047 | 0.04683485 |
| *Mrpl53* | 422.367295 | 0.34530196 | 0.01558959 |
| *Bud31* | 488.82627 | 0.34502248 | 0.02285662 |
| *Selenot* | 2891.26573 | 0.3447606 | 0.00094084 |
| *Echs1* | 905.565763 | 0.34453625 | 0.00230684 |
| *Rpl5* | 4656.53789 | 0.34450738 | 0.02646808 |
| *Acer3* | 268.314213 | 0.3436206 | 0.03160297 |
| *Nr1d2* | 1941.63642 | 0.34294086 | 0.00022375 |
| *Etv1* | 1376.43674 | 0.34291572 | 0.00034827 |
| *Chmp2b* | 595.211662 | 0.34269297 | 0.0092212 |
| *Atp5if1* | 1607.52386 | 0.34261508 | 0.02107446 |
| *S100a16* | 1060.88903 | 0.34257009 | 0.0290849 |
| *Chmp2a* | 920.381086 | 0.3423677 | 0.00551414 |
| *Glrx3* | 1091.94796 | 0.34168105 | 0.00369641 |
| *Ppia* | 11045.0805 | 0.3415795 | 0.00862459 |
| *Tmem128* | 286.049508 | 0.34049699 | 0.03666908 |
| *Nae1* | 522.413133 | 0.34028312 | 0.00541513 |
| *Rps15a* | 2276.10767 | 0.33971866 | 0.02635808 |
| *Ddt* | 444.242207 | 0.33935646 | 0.04669783 |

| | | | |
|---|---|---|---|
| *Gpx1* | 1002.50858 | 0.33782404 | 0.0256611 |
| *Cfl2* | 738.247118 | 0.33703698 | 0.00862459 |
| *Erg28* | 303.519928 | 0.33681122 | 0.01558959 |
| *Mob1a* | 342.004155 | 0.33660928 | 0.01441162 |
| *Eif3m* | 955.345305 | 0.33581809 | 0.00859357 |
| *Pnrc2* | 681.174367 | 0.33538355 | 0.01825661 |
| *Capza2* | 2847.64007 | 0.33472458 | 0.00093949 |
| *D3Ertd751e* | 192.04424 | 0.33431053 | 0.03785501 |
| *Cycs* | 2070.03369 | 0.33340537 | 0.00230684 |
| *Fpgt* | 195.197309 | 0.33291325 | 0.04542842 |
| *Arpc5* | 1146.74606 | 0.33257301 | 0.01782609 |
| *Eef1akmt2* | 203.200849 | 0.33193969 | 0.04858859 |
| *Mrpl50* | 523.673714 | 0.33167029 | 0.04512373 |
| *Ube2b* | 1451.42624 | 0.33157121 | 0.00256791 |
| *Rap1b* | 762.887739 | 0.33081979 | 0.01656208 |
| *Ier3ip1* | 587.351109 | 0.32915081 | 0.02142217 |
| *Hint1* | 1557.20198 | 0.32886744 | 0.0402499 |
| *Atp5pf* | 1786.32452 | 0.32874373 | 0.00313367 |
| *Zfp35* | 214.06583 | 0.32869999 | 0.04264951 |
| *Atp5pd* | 2609.17086 | 0.32839739 | 0.00862459 |
| *Bloc1s2* | 338.068658 | 0.32828166 | 0.01782609 |
| *Blvra* | 212.033553 | 0.32791034 | 0.04324808 |
| *Atp6v1g1* | 1220.86671 | 0.32757083 | 0.00862459 |
| *Btf3l4* | 925.091018 | 0.32730025 | 0.00202836 |
| *Mrpl11* | 460.642233 | 0.32704916 | 0.03479579 |
| *Mrpl46* | 307.549487 | 0.32658675 | 0.04683059 |
| *Snrnp27* | 300.749868 | 0.32645669 | 0.04267558 |
| *Tmem256* | 498.404505 | 0.32623061 | 0.04882284 |
| *Selenow* | 6472.57077 | 0.32622867 | 0.01242449 |
| *Zfp120* | 194.295254 | 0.32576297 | 0.0462101 |
| *Cacybp* | 834.794309 | 0.32554765 | 0.00248676 |
| *Smim20* | 269.882238 | 0.32485305 | 0.0370572 |
| *Supt4a* | 540.957244 | 0.32441412 | 0.03997118 |
| *Zfp938* | 240.662187 | 0.32381487 | 0.03074496 |
| *Nt5c3* | 491.874707 | 0.32358818 | 0.03977528 |
| *Rpl27* | 1545.7976 | 0.32344789 | 0.03230327 |
| *Tomm7* | 673.397108 | 0.32335465 | 0.04127241 |
| *Tvp23b* | 412.131316 | 0.32332193 | 0.02453069 |
| *Aasdhppt* | 272.832677 | 0.32310958 | 0.02285662 |
| *Synpr* | 1689.64251 | 0.32282064 | 0.00200338 |

| | | | |
|---|---|---|---|
| *Golga7* | 1441.42759 | 0.32251309 | 0.00670618 |
| *Kitl* | 496.855907 | 0.3216419 | 0.00705974 |
| *Cibar1* | 494.193142 | 0.32147791 | 0.0261588 |
| *Idi1* | 732.146283 | 0.32114678 | 0.00844262 |
| *Cox14* | 772.644357 | 0.32112796 | 0.02415713 |
| *Lsm1* | 246.706845 | 0.32089769 | 0.03767933 |
| *Nudt19* | 885.611225 | 0.32001817 | 0.04314042 |
| *Tmem68* | 409.782531 | 0.31987154 | 0.02529826 |
| *Cav2* | 423.380059 | 0.31984506 | 0.02616402 |
| *Rpl41* | 3326.49389 | 0.31940521 | 0.04797401 |
| *Pomp* | 1149.41559 | 0.31929266 | 0.00862459 |
| *Eloc* | 1269.97716 | 0.31879703 | 0.02456346 |
| *Anapc11* | 654.266255 | 0.3185567 | 0.00705974 |
| *Rgs5* | 891.657986 | 0.3185538 | 0.00065432 |
| *Btf3* | 1668.78569 | 0.31595779 | 0.00886749 |
| *Iqcb1* | 352.414774 | 0.3157051 | 0.04011688 |
| *Mrps18c* | 366.948526 | 0.31467456 | 0.02325308 |
| *Zfp617* | 415.01615 | 0.31458064 | 0.03445062 |
| *Thoc7* | 680.13105 | 0.31457536 | 0.00952846 |
| *Dram2* | 258.795561 | 0.31427007 | 0.04629228 |
| *Cisd2* | 718.73876 | 0.31383353 | 0.00862459 |
| *Pex13* | 411.878576 | 0.31376104 | 0.02298864 |
| *Yipf5* | 580.968632 | 0.312361 | 0.03076348 |
| *Skic8* | 484.707667 | 0.31235792 | 0.01110346 |
| *Guk1* | 1224.12752 | 0.31090523 | 0.00821264 |
| *Pkia* | 2289.41198 | 0.31087981 | 0.00988317 |
| *Txndc9* | 470.786905 | 0.31079344 | 0.0092212 |
| *mt-Co2* | 53666.4974 | 0.30973943 | 0.01777441 |
| *Eef1b2* | 1821.97943 | 0.30948795 | 0.01661174 |
| *Rpl36al* | 511.437947 | 0.30941763 | 0.0161822 |
| *Psma1* | 858.197791 | 0.30933543 | 0.01321133 |
| *Vamp3* | 448.604726 | 0.30836004 | 0.02857625 |
| *Ndufa8* | 1374.00026 | 0.30827452 | 0.01827321 |
| *Smim7* | 1267.65285 | 0.30822053 | 0.00827754 |
| *Hikeshi* | 385.396236 | 0.30671918 | 0.02897071 |
| *Phospho2* | 340.893489 | 0.30637921 | 0.02453069 |
| *Tmem33* | 1136.69876 | 0.30617865 | 0.00652799 |
| *Brk1* | 1619.5886 | 0.30574017 | 0.03465087 |
| *Ptcd3* | 582.580535 | 0.30530746 | 0.02325308 |
| *Cops4* | 961.762793 | 0.30495901 | 0.0092212 |

| | | | |
|---|---|---|---|
| *Atp5mf* | 1439.3844 | 0.30476068 | 0.04324808 |
| *Lrrc57* | 325.087446 | 0.3046379 | 0.03666908 |
| *Atp6v1f* | 864.346653 | 0.3043134 | 0.0088412 |
| *Crbn* | 952.410439 | 0.3037334 | 0.00862459 |
| *Ndufb8* | 2029.52357 | 0.30348897 | 0.02453069 |
| *H2az2* | 428.60173 | 0.30312218 | 0.03801882 |
| *Higd1a* | 1019.36759 | 0.30310598 | 0.01724057 |
| *Pex7* | 360.379992 | 0.3019668 | 0.02529826 |
| *Uqcrb* | 766.804322 | 0.30191416 | 0.01441162 |
| *Clk1* | 1505.56741 | 0.30146667 | 0.03465087 |
| *Pno1* | 318.317295 | 0.30137607 | 0.03181119 |
| *Lztfl1* | 531.210756 | 0.30087439 | 0.01844832 |
| *Arpp19* | 2710.13912 | 0.30065191 | 0.00705974 |
| *Mrps23* | 475.157628 | 0.30044755 | 0.01476885 |
| *Sub1* | 2009.09466 | 0.30034479 | 0.01302628 |
| *Serp1* | 583.989702 | 0.30001552 | 0.02242778 |
| *mt-Atp6* | 33168.162 | 0.29970862 | 0.01441162 |
| *Atp5f1c* | 2738.47114 | 0.29921367 | 0.04924873 |
| *Polr1d* | 575.446373 | 0.29919679 | 0.01734963 |
| *Tomm22* | 691.564192 | 0.29863891 | 0.0172333 |
| *Sec11c* | 1028.05033 | 0.29820722 | 0.02498746 |
| *Zfand6* | 580.434198 | 0.29719959 | 0.02285662 |
| *Slc25a33* | 665.280092 | 0.29709331 | 0.02545871 |
| *Crot* | 384.615925 | 0.29707907 | 0.03702189 |
| *Mpc1* | 1413.15156 | 0.29616514 | 0.0088412 |
| *Tmem70* | 486.223727 | 0.29584944 | 0.02142217 |
| *Ap4s1* | 410.803382 | 0.29539541 | 0.03814931 |
| *Paip2* | 1674.79092 | 0.29514659 | 0.01476885 |
| *Ptcd2* | 296.42592 | 0.29503519 | 0.04264951 |
| *Cops2* | 987.179392 | 0.29495222 | 0.01537298 |
| *Ran* | 2482.29532 | 0.29449061 | 0.00222097 |
| *Rps11* | 2510.52289 | 0.29298513 | 0.04504846 |
| *Thap12* | 1281.05972 | 0.29225081 | 0.0141086 |
| *Tmed2* | 1348.57366 | 0.29220402 | 0.00862459 |
| *Rps14* | 3224.65547 | 0.29216643 | 0.0479108 |
| *Cd63* | 868.251573 | 0.2921154 | 0.02325308 |
| *Micos10* | 1011.70923 | 0.29190676 | 0.01479725 |
| *Mrpl24* | 466.821185 | 0.29140798 | 0.02236469 |
| *Mpc2* | 1119.66831 | 0.29138836 | 0.03323118 |
| *Pcna* | 346.04723 | 0.29082897 | 0.02079349 |

| | | | |
|---|---|---|---|
| *Cox5b* | 2879.06232 | 0.29035937 | 0.0212338 |
| *Phpt1* | 411.128396 | 0.28978302 | 0.02874252 |
| *Sft2d1* | 342.832705 | 0.28952914 | 0.04394977 |
| *Rpl19* | 4010.55159 | 0.28915374 | 0.01825661 |
| *Olfm3* | 482.436917 | 0.28793034 | 0.01123544 |
| *Nrsn1* | 5207.39379 | 0.28726666 | 0.02656964 |
| *Svip* | 1055.58708 | 0.28696702 | 0.03830159 |
| *Ndufb9* | 2421.53896 | 0.28685488 | 0.02498746 |
| *Lsm6* | 532.401329 | 0.28679013 | 0.01090116 |
| *Ergic2* | 522.067529 | 0.2863405 | 0.01656208 |
| *Rnf7* | 1116.15537 | 0.28613266 | 0.02415697 |
| *Mff* | 2779.74008 | 0.28550963 | 0.03072986 |
| *Rps5* | 2365.02929 | 0.2848731 | 0.03830159 |
| *Nipa2* | 322.957351 | 0.28441522 | 0.04629228 |
| *Ptp4a1* | 1468.50895 | 0.28434157 | 0.01633498 |
| *Mob1b* | 320.403186 | 0.28404609 | 0.03465087 |
| *Hsbp1* | 2352.04133 | 0.28327543 | 0.01782609 |
| *Tcea1* | 1086.11673 | 0.28325654 | 0.0161822 |
| *Ssu72* | 616.792682 | 0.2828016 | 0.01782609 |
| *Rsl24d1* | 603.305486 | 0.28178722 | 0.02034553 |
| *Esd* | 784.729661 | 0.28147197 | 0.04489412 |
| *Dcun1d5* | 329.896533 | 0.28135096 | 0.03294305 |
| *Dnajc15* | 970.958849 | 0.28082649 | 0.02812222 |
| *Timm23* | 881.167482 | 0.28056153 | 0.02176595 |
| *Gria4* | 1688.00595 | 0.28036535 | 0.00859357 |
| *Arrdc3* | 721.65127 | 0.27875927 | 0.0443231 |
| *Psma4* | 748.186432 | 0.27821104 | 0.02236939 |
| *Sdhd* | 1109.24515 | 0.27813758 | 0.04182945 |
| *Map1lc3b* | 3226.13401 | 0.27777582 | 0.01023318 |
| *Uqcrc2* | 2538.39378 | 0.27743348 | 0.00369641 |
| *Jkamp* | 539.648588 | 0.27735109 | 0.0212338 |
| *Srsf7* | 1139.61292 | 0.27661131 | 0.04392684 |
| *Atp5po* | 2352.34484 | 0.27635594 | 0.02903652 |
| *Lin7c* | 780.263653 | 0.27630513 | 0.00248676 |
| *Zfp68* | 431.951006 | 0.27613473 | 0.04011688 |
| *Fgfr1op2* | 1136.76737 | 0.27575394 | 0.01042543 |
| *Ift22* | 579.479912 | 0.27551159 | 0.01844103 |
| *Cggbp1* | 618.648002 | 0.27518123 | 0.01305719 |
| *Ndufab1* | 999.159482 | 0.27460178 | 0.02498746 |
| *Zranb2* | 2045.92624 | 0.27432176 | 0.00359634 |

| | | | |
|---|---|---|---|
| *Mrpl48* | 1075.71071 | 0.27353333 | 0.01915434 |
| *Ppp1cb* | 2976.9788 | 0.27345317 | 0.00067601 |
| *Cdc26* | 408.151285 | 0.27327042 | 0.04519672 |
| *Zc2hc1a* | 571.926294 | 0.27314883 | 0.01827321 |
| *Chchd2* | 3115.04261 | 0.27246947 | 0.04683059 |
| *Vmp1* | 1308.04761 | 0.27228456 | 0.00433875 |
| *Dpy19l4* | 372.867179 | 0.27197485 | 0.0393252 |
| *Gmfb* | 2044.83492 | 0.27081904 | 0.00675485 |
| *Rpl7l1* | 515.645891 | 0.27066042 | 0.03977182 |
| *Atp6v0b* | 2738.91837 | 0.26987421 | 0.02757035 |
| *Pdcd6* | 686.94009 | 0.26911611 | 0.04201752 |
| *Mat2b* | 1376.78153 | 0.26841356 | 0.01827321 |
| *Zcrb1* | 531.277491 | 0.26798303 | 0.02498746 |
| *Ggps1* | 783.517887 | 0.26755333 | 0.01825661 |
| *Zfp260* | 603.045151 | 0.2670305 | 0.02027538 |
| *Ufm1* | 746.57179 | 0.26680199 | 0.03850775 |
| *Chordc1* | 577.942992 | 0.26670559 | 0.01368721 |
| *Ube2d3* | 2807.17373 | 0.26620343 | 0.03479579 |
| *Atp5pb* | 3235.43373 | 0.26613771 | 0.02415697 |
| *Cox6c* | 3369.56719 | 0.26509351 | 0.03302955 |
| *Cct4* | 1210.91579 | 0.26480982 | 0.01402437 |
| *Sdhaf2* | 514.885218 | 0.26329489 | 0.04512373 |
| *Sumo2* | 1422.88986 | 0.26294388 | 0.03767933 |
| *Tsg101* | 626.395416 | 0.26273585 | 0.03766312 |
| *Dynll1* | 2055.17213 | 0.2626799 | 0.04280729 |
| *Rab11a* | 1141.92372 | 0.26226641 | 0.01372536 |
| *Fyttd1* | 943.574341 | 0.26212798 | 0.03496351 |
| *Ssr3* | 1604.54188 | 0.26084777 | 0.00976253 |
| *Txnl1* | 683.247563 | 0.26072678 | 0.01483541 |
| *Cox4i1* | 7742.23055 | 0.26070985 | 0.02498746 |
| *Dnajb9* | 439.280404 | 0.26030964 | 0.04154621 |
| *Jag2* | 453.05683 | -0.2607194 | 0.02673392 |
| *Cers1* | 808.538518 | -0.2609625 | 0.0212338 |
| *Ptprn* | 4562.69444 | -0.2615375 | 0.00859357 |
| *Sorcs2* | 505.910844 | -0.2631528 | 0.03814931 |
| *Ncor2* | 2087.39279 | -0.263197 | 0.00862459 |
| *Col16a1* | 406.586812 | -0.2638923 | 0.03281345 |
| *Mpp2* | 1172.42364 | -0.2649671 | 0.00650312 |
| *Zmiz2* | 2735.76669 | -0.2659478 | 0.01782609 |
| *Fkbp9* | 377.336187 | -0.2671124 | 0.03722466 |

| | | | |
|---|---|---|---|
| *Sema6b* | 1236.44243 | -0.2678871 | 0.0256611 |
| *Vat1* | 510.48499 | -0.2695055 | 0.01934747 |
| *Ehmt2* | 1901.90448 | -0.2695695 | 0.0092212 |
| *Nlgn2* | 3071.62103 | -0.2704507 | 0.00705459 |
| *Tmem130* | 1959.52857 | -0.2715358 | 0.00369641 |
| *Crtac1* | 493.030611 | -0.2715488 | 0.02331741 |
| *Chrna4* | 502.76594 | -0.2725314 | 0.04091326 |
| *Wfs1* | 1005.71785 | -0.2730516 | 0.03332833 |
| *Tmem8b* | 884.172389 | -0.2753159 | 0.02957804 |
| *Sema6c* | 290.909341 | -0.2765339 | 0.04267558 |
| *Zdhhc8* | 973.004926 | -0.277678 | 0.0168916 |
| *Celf5* | 1726.40933 | -0.2778346 | 0.01526955 |
| *Lrfn1* | 474.027486 | -0.2781521 | 0.01838863 |
| *Mbd6* | 604.607478 | -0.2790107 | 0.0163617 |
| *Lrrc47* | 478.935867 | -0.2790532 | 0.04293566 |
| *Lrrc4b* | 1686.75121 | -0.2797944 | 0.00369641 |
| *Wipf3* | 1402.22731 | -0.2802457 | 0.03582992 |
| *Scaf1* | 2132.58294 | -0.2844543 | 0.00380965 |
| *Tmem63a* | 520.996598 | -0.2847775 | 0.01724057 |
| *Grik5* | 1987.20265 | -0.2849303 | 0.01476885 |
| *Chd3* | 5790.30307 | -0.2850933 | 0.00652799 |
| *Myrf* | 875.878282 | -0.2850974 | 0.00859357 |
| *Cacnb1* | 939.642273 | -0.2851451 | 0.01023318 |
| *Hspa2* | 369.660495 | -0.2855759 | 0.02383218 |
| *Vgf* | 1479.20439 | -0.2875766 | 0.0092212 |
| *Git1* | 3810.11334 | -0.2878027 | 0.0168916 |
| *Gaa* | 3778.84564 | -0.2882305 | 0.00093949 |
| *Mast3* | 2338.61247 | -0.2906477 | 0.01795213 |
| *Slc32a1* | 1491.24518 | -0.2918869 | 0.01795213 |
| *Irf2bp1* | 456.414263 | -0.2943563 | 0.04314042 |
| *Rexo1* | 628.993362 | -0.2944944 | 0.00862459 |
| *Pcdh8* | 346.08775 | -0.2949441 | 0.03785501 |
| *Shank3* | 3041.66454 | -0.2960245 | 0.03814931 |
| *Shc2* | 474.822704 | -0.296675 | 0.02176595 |
| *Zfp316* | 332.604468 | -0.2968094 | 0.02812222 |
| *Lmtk3* | 2526.20616 | -0.2970051 | 0.00087596 |
| *Tnk2* | 2762.05202 | -0.2973896 | 0.00093949 |
| *Rgma* | 684.917266 | -0.2994165 | 0.0046996 |
| *Shisa7* | 1081.41525 | -0.3007019 | 0.0046996 |
| *Slc6a11* | 2952.28572 | -0.3010921 | 0.0011208 |

| | | | |
|---|---|---|---|
| *Tmem94* | 825.062868 | -0.301188 | 0.00559951 |
| *Cbarp* | 2864.984 | -0.3012967 | 0.00096569 |
| *Foxg1* | 724.158055 | -0.3016019 | 0.02285662 |
| *Polr1a* | 394.976351 | -0.3026821 | 0.01302628 |
| *Sez6* | 2020.82716 | -0.3056765 | 0.00719293 |
| *Cacng4* | 468.048684 | -0.3077436 | 0.01123544 |
| *Adgrb2* | 3474.74825 | -0.3081928 | 0.0088412 |
| *Ccdc92b* | 376.373071 | -0.3084327 | 0.03406097 |
| *Phf2* | 730.816231 | -0.3093277 | 0.00369641 |
| *Nptx2* | 234.859152 | -0.312525 | 0.04981141 |
| *Kcnj4* | 604.425963 | -0.3160803 | 0.04267558 |
| *Nrp2* | 384.109487 | -0.3194527 | 0.01090116 |
| *Ssbp4* | 734.429567 | -0.3216692 | 0.00202836 |
| *Atn1* | 1911.86384 | -0.3230634 | 0.00128245 |
| *Zfp319* | 282.248826 | -0.3247405 | 0.02236469 |
| *Rgs9* | 548.842026 | -0.3280885 | 0.02782402 |
| *Adcy5* | 3193.8425 | -0.3323958 | 0.03160297 |
| *Grip2* | 408.08663 | -0.3340322 | 0.00859357 |
| *Tox2* | 338.58464 | -0.3362557 | 0.02732814 |
| *Limk1* | 1027.25099 | -0.3372732 | 0.00088785 |
| *Rasal1* | 632.856454 | -0.3374163 | 0.00924179 |
| *Sema5b* | 206.905574 | -0.3409799 | 0.0364276 |
| *Cpne5* | 668.44757 | -0.3485985 | 0.0168916 |
| *Cactin* | 272.15381 | -0.349578 | 0.02236469 |
| *Cacng8* | 499.239142 | -0.3544479 | 0.0092212 |
| *Slit1* | 787.597902 | -0.3568099 | 0.00136166 |
| *Dscaml1* | 549.599862 | -0.3731594 | 0.00131444 |
| *Cacna1h* | 816.660673 | -0.4001402 | 0.00037146 |
| *Hcn4* | 127.625755 | -0.4075674 | 0.04860764 |
| *Baiap3* | 714.167006 | -0.410072 | 0.01463513 |
| *Pcdhgc4* | 320.631477 | -0.4202589 | 0.02990791 |
| *Zfp628* | 126.497466 | -0.4520289 | 0.04542842 |
| *Fzd2* | 120.735636 | -0.4891012 | 0.02790403 |
